# Feature-reweighted representational similarity analysis: A method for improving the fit between computational models, brains, and behavior

**DOI:** 10.1101/2021.09.27.462005

**Authors:** Philipp Kaniuth, Martin N. Hebart

**Affiliations:** Vision and Computational Cognition Group, Max Planck Institute for Human Cognitive and Brain Sciences, Stephanstrasse 1A, Leipzig, Germany

**Author notes:** Corresponding authors. ( —).

**Keywords:** Representational similarity analysis, multivariate pattern analysis, functional MRI, MEG, behavior, deep neural networks, noise ceilings

## Abstract

Representational Similarity Analysis (RSA) has emerged as a popular method for relating representational spaces from human brain activity, behavioral data, and computational models. RSA is based on the comparison of representational (dis-)similarity matrices (RDM or RSM), which characterize the pairwise (dis-)similarities of all conditions across all features (e.g. fMRI voxels or units of a model). However, classical RSA treats each feature as equally important. This ‘equal weights’ assumption contrasts with the flexibility of multivariate decoding, which reweights individual features for predicting a target variable. As a consequence, classical RSA may lead researchers to underestimate the correspondence between a model and a brain region and, in case of model comparison, may lead them to select an inferior model. The aim of this work is twofold: First, we sought to broadly test feature-reweighted RSA (FR-RSA) applied to computational models and reveal the extent to which reweighting model features improves RSM correspondence and affects model selection. Previous work suggested that reweighting can improve model selection in RSA but it has remained unclear to what extent these results generalize across datasets and data modalities. To draw more general conclusions, we utilized a range of publicly available datasets and three popular deep neural networks (DNNs). Second, we propose voxel-reweighted RSA, a novel use case of FR-RSA that reweights fMRI voxels, mirroring the rationale of multivariate decoding of optimally combining voxel activity patterns. We found that reweighting individual model units markedly improved the fit between model RSMs and target RSMs derived from several fMRI and behavioral datasets and affected model selection, highlighting the importance of considering FR-RSA. For voxel-reweighted RSA, improvements in RSM correspondence were even more pronounced, demonstrating the utility of this novel approach. We additionally show that classical noise ceilings can be exceeded when FR-RSA is applied and propose an updated approach for their computation. Taken together, our results broadly validate the use of FR-RSA for improving the fit between computational models, brain, and behavioral data, possibly allowing us to better adjudicate between competing computational models. Further, our results suggest that FR-RSA applied to brain measurement channels could become an important new method to assess the correspondence between representational spaces.

## 1. Introduction

A core aim of cognitive neuroscience is to reveal the nature of our neural representations and determine their role in shaping cognition and overt behavior. Central to this aim are comparisons of representations measured in brain activity data (e.g. fMRI, MEG) with representations derived from computational models or behavior. A powerful framework for such comparisons is offered through representational similarity analysis (RSA). RSA abstracts away from the measurement level (e.g. voxels, sensors) to the level of representational (dis-)similarities, allowing for direct comparisons across modalities, species, models, and behavior (Kriegeskorte et al., 2008a; Kriegeskorte and Kievit, 2013). By characterizing representations as (dis-)similarities of activity patterns, RSA has become a central tool for multivariate pattern analysis, complementing multivariate decoding (Haynes and Rees, 2006; Hebart and Baker, 2018) and other methods operating at the level of multivariate activity patterns (Haxby et al., 2014; Diedrichsen et al., 2018). RSA is not only useful for identifying the presence of a representational correspondence between modalities; it also provides a simple yet effective approach for comparing computational models with behavioral and neuroimaging data.

At the heart of RSA lies the computation of representational (dis-)similarity matrices (RDMs or RSMs), which characterize the (dis-)similarity of all pairs of conditions (e.g. visual stimuli) across all features (e.g. measurement channels, units of a computational model). While in recent years a lot of focus has been placed on improving the reliability of RDMs (Walther et al., 2016; Charest et al., 2018) and identifying the most appropriate dissimilarity measure (Walther et al., 2016; Bobadilla-Suarez et al., 2020; Ramírez et al., 2020), much less emphasis has been placed on the contribution of individual features in the computation of representational (dis-)similarities. In fact, most RSA approaches assume that each feature is of equal importance and will thus contribute equally to the final (dis-)similarity estimate. This ’equal weights’ assumption is at odds with the idea that for a given comparison of RSMs, some features may carry more information than others. This has several important consequences. First, for computational models, classical RSA may underestimate the correspondence between the model and a given brain region, even though the model may already contain the relevant representational space for capturing the brain response. This may lead not only to suboptimal model performance, but may also affect different models to different degrees, which in case of model comparisons may lead to the selection of an inferior model (Khaligh-Razavi and Kriegeskorte, 2014; Peterson et al., 2016; Jozwik et al., 2017; Storrs et al., 2021). Second, for brain data, classical RSA may overemphasize the importance of individual brain measurement channels, treating noisy channels (e.g. voxels) as equally important as channels that carry signal. This contrasts with the intuition used in multivariate linear decoding, where each voxel’s importance is reweighted according to the contribution to the final classification task (Figure 1a). Surprisingly, reweighting of individual brain measurement channels is not routinely applied to the measurement of representational similarities. This suggests large untapped potential for improving the representational correspondence between computational models, brain activity, and behavior (Figure 1b,c).

**Figure 1:**
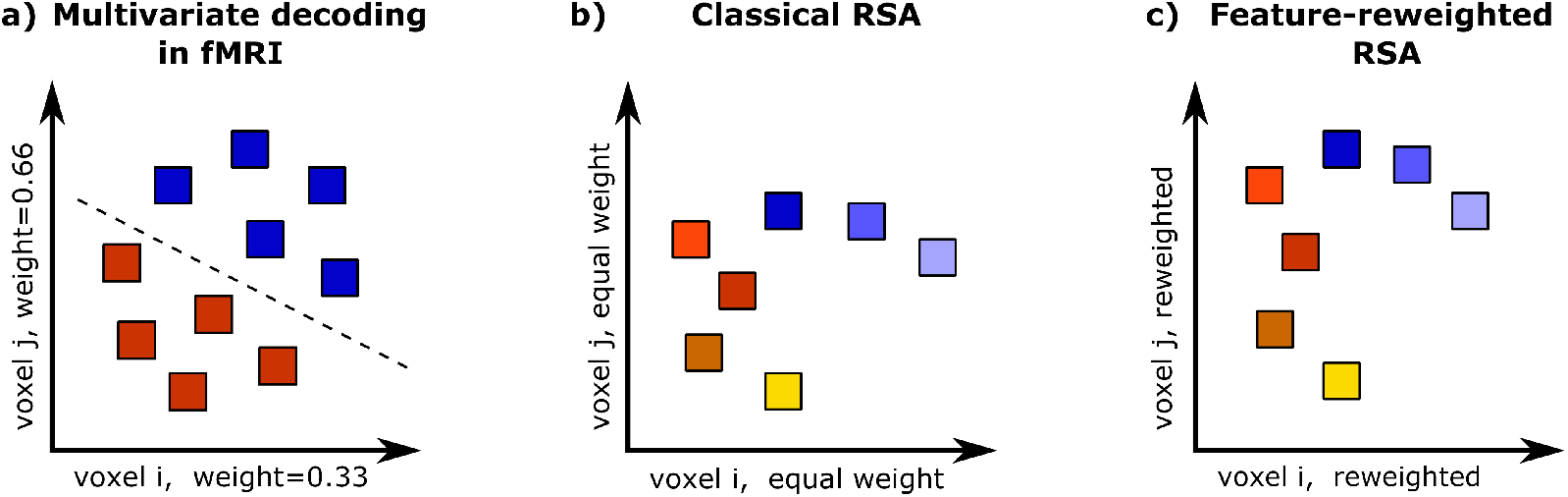
Parallel between multivariate decoding and feature-reweighted RSA (FR-RSA). (a) Multivariate decoding in fMRI can be described as a linear reweighting of features (e.g. voxels) to optimize the linear readout of a binary target variable (e.g. stimulus category). Voxel weights reflect their importance for the final classification objective. (b) In classical RSA, no reweighting is applied, which assumes that all voxels are equally important. Relative to multivariate decoding, this underestimates the linear information contained in multivoxel activation patterns. (c) FR-RSA utilizes linear feature-reweighting in order to best reflect pairwise (dis-)similarity relative to a target RDM or RSM, thereby improving the use of multivariate information present in the data.

The aim of the present study is twofold. First, for the reweighting of computational model units, we seek to broadly validate the degree to which feature-reweighted RSA (FR-RSA) can act as a general-purpose method to relate representational spaces of models to those of brain and behavior. To achieve this aim, we systematically apply FR-RSA to representations from deep neural networks (DNN) on the one hand and relate them to diverse publicly available neuroimaging and behavioral datasets on the other hand. Second, for the reweighting of brain measurement channels, we demonstrate the broad applicability of FR-RSA applied to fMRI data—*voxel-reweighted* RSA—for improving the correspondence between representational similarities derived from the brain and from models or behavior. Previewing our results, we find that reweighting units of a DNN reliably improves the fit between model, brain, and behavioral RSMs and indeed affects which DNN is selected as the best model of brain activity (Storrs et al., 2021). This generalizes the utility of FR-RSA to a broad set of neural network models, brain imaging methods, behavior, and stimuli. Further, when reweighting is applied to fMRI voxels, our results demonstrate consistent and pronounced improvements of RSM correspondence. This suggests that feature reweighting applied at the level of brain measurements may act as a general-purpose method for improving the representational correspondence between brains, models, and behavior. To facilitate future use of this method, we provide a toolbox to run FR-RSA in Python (https://github.com/ViCCo-Group/frrsa), with recommendations regarding implementational choices.

## 2. Methods

### 2.1. Datasets and computational models

We sought to evaluate the general applicability of feature-reweighted RSA (FR-RSA), both when (1) reweighting individual units of a computational model, as has been done previously with similar approaches (e.g. Peterson et al., 2016; Jozwik et al., 2017; Storrs et al., 2021) and (2) when reweighting measurement channels of brain data, an approach which to our knowledge has not been carried out before. To this end, we used datasets from several published studies in which participants had been exposed to a range of object images (Mur et al., 2013; Cichy et al., 2014, 2016; Bankson et al., 2018; Cichy et al., 2019). The datasets are centered around four sets of natural object images and reflect a combination of functional MRI data, magnetoencephalography data, and behavioral similarity judgments. In addition, for the object images, we extracted neural network activations as computational models. Together, this makes these datasets well suited for evaluating FR-RSA across a wide range of possible analyses. One of the published studies (Bankson et al., 2018) used a twin set of 84 natural object images, which were tested in separate sets of participants and which we thus treated as two separate datasets. Another image set (Kriegeskorte et al., 2008b; Mur et al., 2013; Cichy et al., 2014) consisted of 92 images of human and non-human faces and bodies, as well as natural and artificial objects. Finally, another image set (Cichy et al., 2016, 2019) consisted of 118 natural images. All datasets used in this work were part of studies with approval by their respective local ethics committee. Details regarding which kinds of data were available for which image set, as well as the task carried out by participants, can be found in Table 1.

**Table 1:** Overview over the datasets used in this study. For each image set, data from several measurement modalities were available. For fMRI, we focused on data from early visual cortex (EVC) and higher visual cortex (HVC). The number of participants, n, for every measurement modality is stated in parentheses. Note that not all available datasets from these image sets were used in all analyses, given the exceedingly large number of possible comparisons.

| Image Set | Type of Images | Experimental Task | MEG (n) | fMRI ROI (n) | MEG Type (channels) | Behavior (n) | Reference |
| --- | --- | --- | --- | --- | --- | --- | --- |
| 84 (Set 1) | natural images cropped and placed on gray background | oddball detection | - | - | - | single arrangement similarity (n = 16) | Bankson et al. (2018) |
| 84 (Set 2) | natural images cropped and placed on gray background | oddball detection | - | - | - | single arrangement similarity (n = 16) | Bankson et al. (2018) |
| 92 | human and non-human faces and bodies as well as natural and artificial objects cropped and placed on gray background | MEG: button press when paperclip shown<br>fMRI: button press when null trial | Yes (16) | Early visual cortex (n = 15)<br>Higher visual cortex (n = 15) | 204 planar gradiometers,<br>102 magnetometers | multiple arrangement similarity (n = 16) | Mur et al. (2013); Cichy et al. (2014) |
| 118 | diverse natural images, on natural background | MEG: button press when paperclip shown<br>fMRI: button press when null trial | Yes (15) | Early visual cortex (n = 15)<br>Higher visual cortex (n = 15) | 204 planar gradiometers,<br>102 magnetometers | multiple arrangement similarity (n=20) | Cichy et al. (2016, 2019) |

#### 2.1.1. fMRI data

For the fMRI data associated with two of the image sets (92 and 118), we used voxel-wise beta estimates for each object. These were provided with the publicly available datasets and had been estimated by applying a general linear model to the preprocessed data. The original studies used a Siemens 3T Trio scanner with a 32-channels head coil and, for the 92 image set, acquired 192 volumes for each participant (gradient-echo EPI sequence: TR = 2,000 ms, TE = 31 ms, flip angle = 80°, FOV read = 192 mm, FOV phase = 100%, ascending acquisition, gap = 10%, resolution = 2 mm isotropic, slices = 25) or, for the 118 image set, 648 volumes for each participant (gradient-echo EPI sequence: TR = 750 ms, TE = 30 ms, flip angle = 61°, FOV read = 192 mm, FOV phase = 100%, ascending acquisition, slice gap = 20%, resolution = 3 mm^3^, slices = 33) (for methodological details, see Cichy et al., 2014, 2016). For simplicity, we focused on early visual cortex (EVC) and higher visual cortex (HVC) as regions of interest. Since data were provided in MNI space only, EVC and HVC were defined using anatomical criteria, based on a projection of the Glasser atlas to MNI space (Glasser et al., 2016). For EVC, we used a mask of areas V1, V2, and V3.

For HVC, we used a mask consisting of areas V8 (VO1), PIT, VVC, FFC, VMV1-3, PHA1-3, TF, and TE2p. For each participant and area, a conservative preselection of voxels was conducted by selecting only the most strongly activated 250 voxels (post-hoc analyses including all voxels showed qualitatively similar yet overall slightly weaker results, results not shown).

#### 2.1.2. MEG data

While human MEG data were available for all image sets, due to the extensive number of possible comparisons, we focused on MEG data for image sets 92 and 118. Both MEG datasets had been acquired with 306 channels at a sampling rate of 1,000 Hz. Data were filtered between 0.03 and 330 Hz and were baseline corrected (for methodological details, see Cichy et al., 2014, 2016). For image set 92, MEG signals were extracted for each trial for 100ms before and 1,200ms after stimulus presentation, resulting in 1,301 samples in total. Across all measurement channels, this yielded a data matrix of size 306 × 92 for every time point and participant. For the image set 118, MEG signals were extracted for 100ms before and 1,000ms after stimulus presentation, resulting in 1,101 samples in total. This yielded a data matrix of size 306 × 118 for every time point and participant.

#### 2.1.3. Behavioral data

The behavioral data of all image sets included in this study were sampled using either the single (Hout et al., 2013) or multiple object arrangement method (Kriegeskorte and Mur, 2012). In those tasks, participants are required to arrange objects in a circular arena according to the perceived dissimilarity between images by dragging-and-dropping them to different locations within the arena. Dissimilar images are positioned further away from each other, while similar images are positioned closer to each other. Importantly for later analyses, this method directly produces fully-sampled RDMs, rather than yielding feature vectors that are then converted into RDMs. The single arrangement method was used for both image sets with 84 images, while the multiple arrangement method was used for image sets 92 and 118. Further details regarding the specifics of behavioral data acquisition can be found in the original studies (Mur et al., 2013; Bankson et al., 2018; Cichy et al., 2019).

#### 2.1.4. Layer activations from chosen deep neural networks

We chose three popular DNN architectures for our investigation: AlexNet (Krizhevsky et al., 2012), VGG-16 (Simonyan and Zisserman, 2015), and ResNet-50 (He et al., 2016). We used versions of the DNNs that had been pretrained on the 1,000 object classes used in the ImageNet Large-Scale Visual Recognition Competition (ILSVRC, Russakovsky et al., 2015), implemented in the Matlab toolbox MatConvNet (Vedaldi and Lenc, 2015). For each DNN, we extracted activity patterns for each image for a subset of DNN layers. For AlexNet, we selected all five pooling layers and the first two fully-connected layers, resulting in seven layers in total. From early to late layers, these layers had 290400, 186624, 64896, 64896, 43264, 4096, and 4096 units, respectively. Similarly, for VGG-16, all five pooling layers and the first two fully connected layers were chosen. From early to late layers, VGG-16’s layers had 802816, 401408, 200704, 100352, 25088, 4096, and 4096 units, respectively. For ResNet-50, layers conv1 (802816 units), res2b (802816 units), res3b (401408 units), res3d (401408 units), res4c (200704 units), res4f (200704 units), and res5c (100352 units) were chosen as roughly corresponding to the chosen layers in AlexNet and VGG-16 in terms of network depth. Subsequently, we will refer to these layers as layers 1 to 7. For each layer, we concatenated all units of all feature maps into one long vector. This yielded one activity pattern per stimulus per layer. No dimensionality reduction was conducted for any DNN layer.

### 2.2. Constructing classical representational similarity matrices

For the MEG and behavioral data we used the RDMs as provided by the original studies and transformed them into RSMs. For the MEG data, original RDMs consisted of pairwise linear support vector machine classification accuracies, where higher accuracies are supposed to reflect better linear separability and thus greater dissimilarity (Cichy et al., 2014, 2016). For the DNN and fMRI activity patterns, when constructing full RSMs, we computed similarities as the Pearson’s correlation coefficient. Behavioral RDMs were chosen as the final result from the object arrangement task (see Behavioral data) and transformed into RSMs.

### 2.3. Reweighting features of a given modality to predict similarities in another modality

#### 2.3.1. Overview over the algorithm

The key differences between classical RSA and FR-RSA are illustrated in Figure 2. In classical RSA, individual cells in one RSM reflect the overall similarity of two activity patterns for conditions *x* and *y* (e.g. DNN layer activations for two object images). The resulting RSM, henceforth called the “predicting RSM”, is then related to another RSM, henceforth called the “target RSM”. In contrast, the rationale of feature-reweighted RSA is that individual RSM cells are treated as a linear combination of univariate, feature-specific similarities. The resulting predicting RSM can thus be conceptualized as a linear combination of feature-specific RSMs (Figure 2b). The aim of FR-RSA is then to learn weights that allow the optimal combination of these feature-specific predicting RSMs in a way that maximizes their correspondence with the target RSM (note that the same reasoning holds for RDMs). In our implementation, this is realized using L2-regularized multiple linear regression in a cross-validation framework.

**Figure 2:**
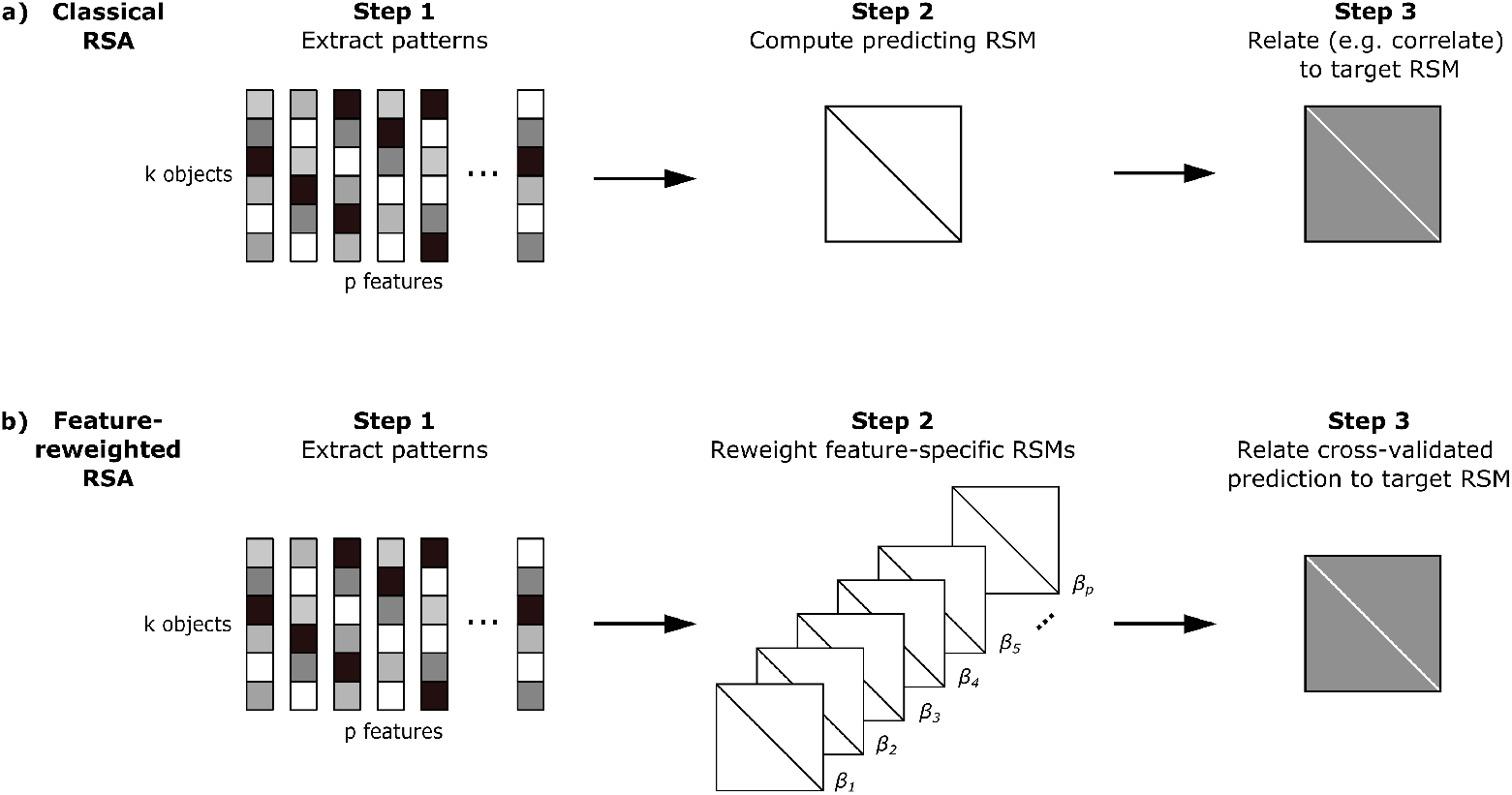
Comparison of classical and feature-reweighted RSA (FR-RSA). **(a)** In classical RSA, the predicting RSM is computed across all features before it is related to the target RSM. **(b)** In FR-RSA, for the predicting RSM, one RSM for each feature is computed. Each feature’s RSM receives its own *β* weight to optimally predict the target RSM using regularized linear regression. All reweighted feature RSMs are then combined and related to the target RSM.

#### 2.3.2. Rationale of the statistical model

More specifically, in classical RSA with Pearson correlation as the similarity measure, an RSM cell is quantified by:

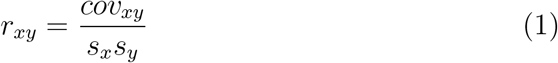

with *x* and *y* referring to the current pair of objects and *s_x_* and *s_y_* referring to the standard deviation of the respective object’s activity pattern. Since the covariance reflects the centered dot product and the correlation coefficient the scaled covariance, it is possible to alternatively z-transform each object pattern, which then reduces the correlation coefficient to the product of the object pairs’ feature values, summed across all features *i*, with a constant scaling factor *p* in the denominator denoting the number of features:

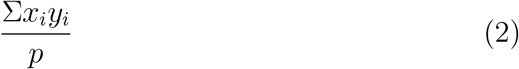

In feature-reweighted RSA, this formula simply translates to:

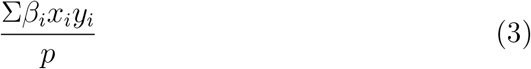

For a given feature *i*, across all object pairs, the same *β* weight is learned. This is equivalent to learning a linear combination of weighted univariate feature-specific RSMs, as illustrated in Figure 2b. Thus, to accurately combine these features and easily fit feature-specific weights, each pattern is z-transformed when using Pearson correlation as the similarity measure. Note that the constant *p* can be ignored as it affects the scaling of all *β* weights equally.

Therefore, the predicting RSM in FR-RSA is a linear combination of feature-specific RSMs, in which each feature-specific RSM receives its own *β* weight. Identifying this weight can be formulated as a multiple regression problem, in which each feature-specific RSM acts as an individual predictor, each with its own unique weight. For this multiple regression model, each feature-specific RSM is flattened so that only its unique upper (or lower) triangular part is used, since each RSM is symmetric along its diagonal.

There are two potential issues with this multiple regression model. First, it can contain a very large number of predictors, which is given by the number of features making up the predicting RSM. Second, given possible redundancy across features, these predictors may exhibit high covariance (e.g. feature-specific RSMs for a number of neighboring voxels). To counteract possible collinearity resulting from these issues, we added a regularization term to the objective function. Since we did not aim at selecting specific features but rather to leverage the complete range of information present in the data, we opted for an L2 regularization, that is, ridge regression (Hoerl and Kennard, 1970), which also offers a closed-form solution. For that purpose we used fractional ridge regression (Rokem and Kay, 2020), since it allows automatic evaluation of the entire range of possible hyperparameters.

Finally, to avoid overfitting, we cross-validated the multiple linear ridge regression model and also conducted a nested cross-validation to establish the best regularization parameter for each cross-validation iteration. The next paragraph provides a step-by-step run-through of the algorithm.

#### 2.3.3. The full sequence of feature-reweighted RSA

1. Data from two measurement modalities (e.g. activity patterns from a specific DNN’s layer and fMRI voxel activities from a pre-defined ROI) are selected. One of the modalities is declared the predicting dataset for which feature-reweighted RSMs are computed. The other modality is the target dataset for which the full RSM is explained by the predicting dataset. In matrix form, the predicting dataset is provided as a *p* × *k* matrix, while the target dataset reflects a full *k* × *k* RSM, with *p* referring to the number of measurement channels and *k* to the number of conditions (see Diedrichsen and Kriegeskorte, 2017).
2. Activity patterns for all conditions (e.g. images) of the predicting dataset are z-transformed (exploratory results when leaving out this step can be found in Supplemental Figure S1).
3. Data are split randomly into five folds for the outer cross-validation. Importantly, data are split along the condition axis, so that the outer training and test set contain non-overlapping sets of condition pairs (different from Peterson et al., 2016, but similar to Jozwik et al., 2017). Other cross-validation schemes are possible but we confirmed with post-hoc analyses that 5-fold cross-validation reflects a good trade-off between speed and accuracy.
4. For a given outer cross-validation iteration, the outer training set is again split repeatedly into five folds, yielding an inner training and inner test set for the inner cross-validation.
5. With the inner cross-validation, the best hyperparameter for ridge regression is estimated. For ridge regression we used fractional ridge regression (Rokem and Kay, 2020). Hence, the hyperparameter to be optimized is the fraction between ordinary least squares and L2-regularized regression coefficients.
6. Once the best hyperparameter for the current outer cross-validation iteration has been established, ridge regression is estimated on the outer training set.
7. The fitted statistical model returns reweighted similarities for the predicting RSM on the test set. Predicted values that are out of range (e.g. predicted correlation coefficients smaller than −1 or larger than 1) are clipped to the nearest permissible value.
8. Finally, predictions are correlated with the respective similarities of the target RSM using Pearson’s *r* to evaluate their fit.

Note that in our implementation, the outer 5-fold cross-validation was repeated ten times and the inner 5-fold cross-validation five times, using different random splits in each iteration. Hence, for a single analysis, 50 outer ridge regression models were fitted. For each of these models, Pearson’s *r* between the reweighted predicted and the respective target similarities was derived, so that all 50 Fisher’s *z*-transformed Pearson’s *r*s were averaged across outer cross-validation folds.

#### 2.3.4. Reweighting analyses conducted in this study

The reweighting analyses we carried out can roughly be divided into two kinds: (1) feature-reweighted RSA applied to DNNs, where activations are reweighted to predict individual participant’s fMRI or behavioral RSMs, and (2) voxel-reweighted RSA, where individual participant’s fMRI activity patterns are reweighted to predict group-averaged behavioral RSMs, DNN RSMs, or group-averaged MEG RSMs (that is, applying reweighting to MEG-fMRI fusion, see Cichy et al., 2014).

Thus, for every reweighting analysis conducted, each participant received one overall score that indicates the correlation between the reweighted predicting RSM and the target RSM. These scores were used for further statistical analyses (similar to Storrs et al., 2021) and compared to classical RSA. Analyses reweighting DNN units were conducted for all image sets, whereas analyses reweighting fMRI voxels were conducted only for the image sets with 92 and 118 images. Analyses involving fMRI were conducted separately for both EVC and HVC ROIs. Analyses involving MEG were conducted separately for every time point. For further statistical analyses and graphical presentation of the results, we averaged every ten samples for results pertaining to MEG data, yielding MEG results for 130 and 110 samples, respectively. Across image sets and use cases, on the group level, this resulted in a total of 736 comparisons of classical and feature-reweighted RSA, reflecting different combinations of a predicting and target RSM. Due to the large number of individual results, we only discuss a selection of them in detail in the main text. For a full overview of all results please refer to the figures. Please note that, in contrast to some prior work (e.g. Jozwik et al., 2017; Storrs et al., 2021), we did not impose a non-negativity constraint on the *β* weights for the main set of analyses (see Supplemental Figure S2 for a subset of analyses with non-negativity constraint). All result files and analysis scripts pertaining to this study are available via an OSF repository (https://osf.io/8weum/). The toolbox to run FR-RSA in Python is available via GitHub (https://github.com/ViCCo-Group/frrsa).

### 2.4. Statistical analyses

#### 2.4.1. Assessing the strength of RSM correspondence

To determine the statistical significance of the correlation between two RSMs at the group-level, we conducted one-sided Wilcoxon signed-rank tests, comparing participants’ rank-transformed correlation values against zero (Nili et al., 2014). Similarly, to test whether RSMs derived from two different computational models are related, to varying degrees, to an RSM derived from either human brain activity or human behavior, we performed twosided Wilcoxon signed-rank tests. We corrected for multiple comparisons by controlling the expected false discovery rate at 0.05 (Benjamini and Hochberg, 1995).

### 2.5. Estimating noise ceilings

Noise ceilings provide an estimate of the best performance any model can achieve given the noise in the data. As is common in RSA, the upper noise ceiling is estimated as the mean correlation between the group-average RSM and each participant-specific RSM. The lower noise ceiling is estimated as the mean correlation between the group-average RSM and each participant-specific RSM while iteratively excluding a given participant from the group-average (Nili et al., 2014).

#### 2.5.1. Estimating reweighted noise ceilings

When conducting feature-reweighted RSA, geometrically speaking, successful reweighting will move the model’s RSM closer to each individual participant’s RSM (or vice versa when reweighting voxels) (see Figure 3). However, at the same time, the position of the group-average RSM relative to the individual participant’s RSMs, which is used for the estimation of the noise ceiling, remains unchanged. As a consequence, for feature-reweighted RSA, classical noise ceilings underestimate the best possible performance any model can achieve since they themselves do not take reweighting into account. This can lead to results that appear to approach or even exceed the noise ceiling while in actual fact this comparison is no longer valid. As a remedy, we propose to also apply reweighting to noise ceilings to get an estimate of the best performance any model can achieve given the noise in the data *and* given that reweighting has been applied. To obtain valid noise ceiling estimates in the context of feature reweighting, for the reweighted upper noise ceiling, we applied reweighting to each participant-specific RSM to optimally predict the group-average RSM and averaged the resulting correlations. For the lower noise ceiling, we reweighted each participant-specific RSM to optimally predict the group-average RSM from which the current participant-specific RSM was left out, again averaging the resulting RSM correlations.

**Figure 3:**
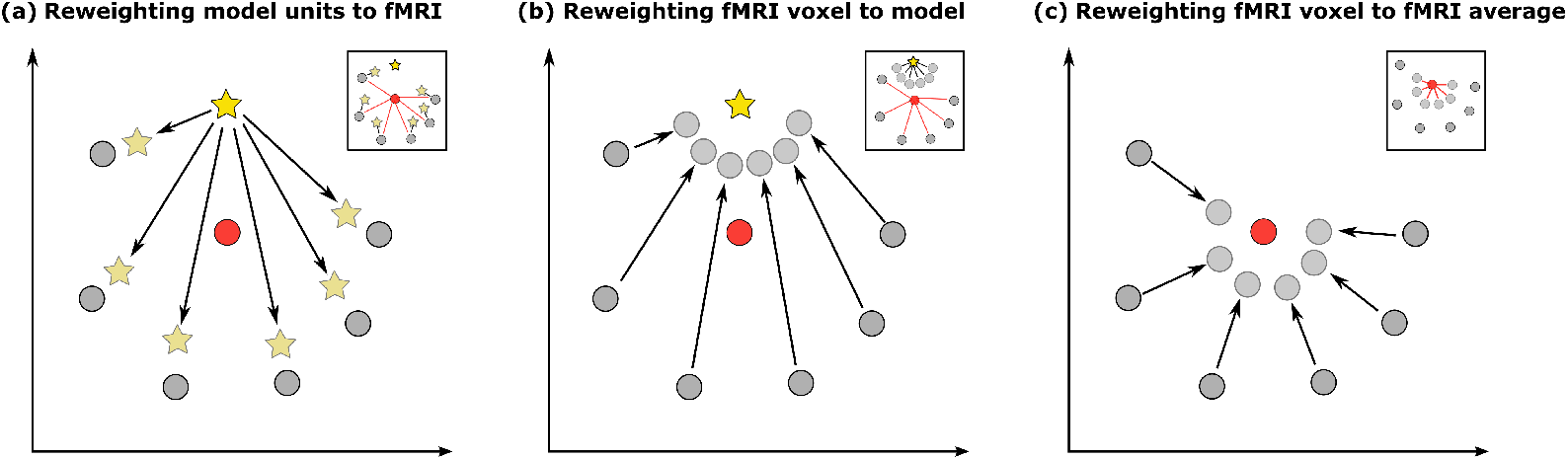
Geometrical reasoning for reweighted noise ceiling. In classical RSA, noise ceilings are calculated based on individual participant’s RSMs (gray circles) and their respective group average (red circle), to get an estimate of the best performance a given model RSM (yellow star) could achieve. **(a)** When a given model’s units are reweighted, the model iteratively moves closer to each participant’s fMRI RSM. Here, the reweighted model may be closer to each participant than the group-average is (small panel). **(b)** Similarly, when reweighting is applied to individual participant’s fMRI voxels, their RSMs move closer towards the model RSM. That way, they can then be closer to the model than the non-reweighted ones are to their mean (small panel). However, when calculating noise ceilings based on the non-reweighted participant RSMs and their respective group average, reweighting is not taken into account. **(c)** Therefore, in either case, noise ceilings should be calculated based on individual participant’s RSMs that have been reweighted to best predict the group-average RSM.

#### 2.5.2. Statistical significance of model relative to noise ceiling

Each correlation between RSMs was tested regarding whether it was significantly below any of the lower noise ceilings, using uncorrected one-sided Wilcoxon signed-rank tests. Not controlling the false discovery rate works against finding a non-significant difference and is therefore a more conservative procedure (Storrs et al., 2021). Note that when a behavioral RSM is the target variable, then estimating reweighted noise ceilings is not possible, given that feature-reweighting cannot be applied without the presence of features (see Behavioral data). In this case, reweighted noise ceilings are omitted.

## 3. Results

### 3.1. Feature-reweighting on simulated data

Prior to conducting feature-reweighted RSA (FR-RSA) on empirical data, we carried out a simulation in which we tested the degree to which FR-RSA is able to (1) improve the correspondence between a model RSM and a target RSM and (2) identify the superior model from a set of competing models. While for empirical data it is not possible to know in advance which is the truly superior model, a simulation can provide a proof of concept for testing whether FR-RSA is able to recover the original ground truth best model and thus improve model selection. To this end, we defined a ground truth representational space *G* with a predefined covariance structure (Figure 4). Next, we created a target RSM that was based on features with the same covariance as *G*, plus multivariate Gaussian noise on these features. Finally, we created two models which we designed in a way that Model 1 was the superior model, while Model 2 was the inferior model.

**Figure 4:**
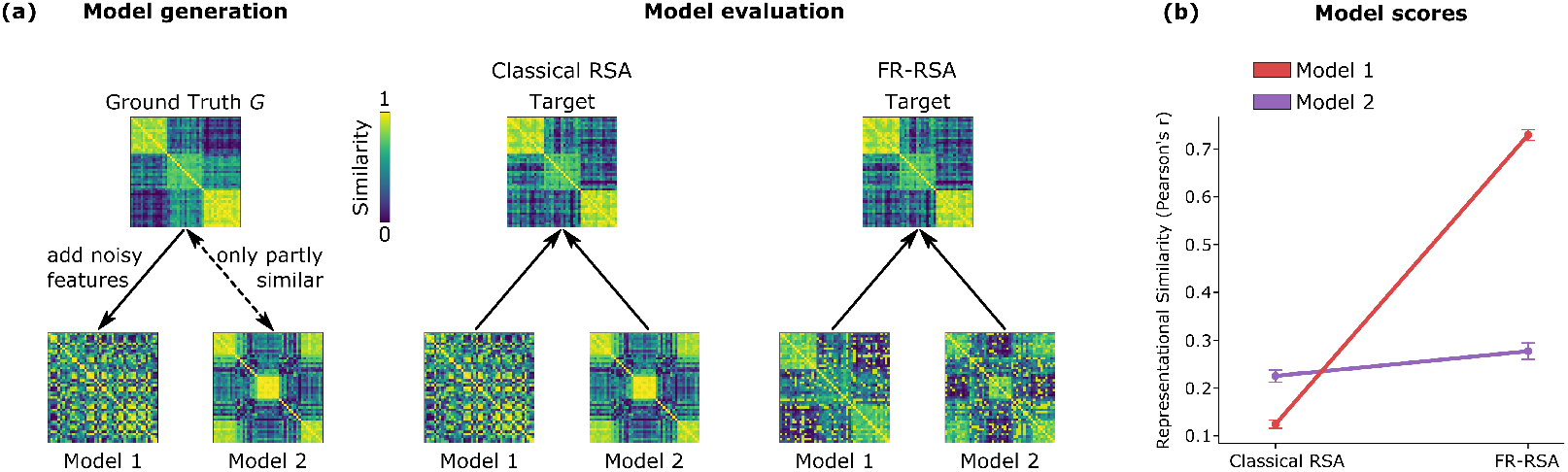
A simulation of the effect of FR-RSA with a known ground truth representational space. **(a) Simulation scheme.** We designed a ground truth representational space, G, based on which we generated two models. Model 1 was designed to be the superior model, which contained the ground truth representational space but was obscured by the addition of a set of non-informative features. Model 2 was designed to be the inferior model by only partially matching ground truth, without the addition of non-informative features. For model evaluation, we then compared both models to a simulated target RSM, which was derived from our ground truth matrix. **(b) Results of the simulation across simulation iterations.** For classical RSA, overall performance was poor, with the inferior Model 2 exhibiting a higher representational correspondence than the superior Model 1. FR-RSA led to a strong improvement in model performance for Model 1, with a much weaker improvement for Model 2, highlighting the potential of FR-RSA to improve representational correspondence while recovering the ground truth representational space.

Model 1 was based on noisy features with the same covariance as G, akin to the creation of the target RSM. However, we added irrelevant features based on pure noise. In contrast, Model 2 was based only partially on the ground truth representational space *G* but without the addition of irrelevant features. Therefore, Model 1 contained the complete relevant subspace defined by *G* but the final representational space was obscured by the addition of irrelevant features, while for Model 2, even its original representational space without the addition of noise would only show a decent correspondence with the target RSM, making it an inferior model as compared to Model 1. We then repeated this simulation 100 times. Having created two competing models and a target RSM, we next conducted classical RSA and FR-RSA on both models, for all 100 simulation iterations. The results of the simulation scheme are presented in Figure 4. Before reweighting, overall performance of both models was poor, with a better fit of Model 2 than Model 1, demonstrating that, in this simulation scheme, classical RSA favored the inferior model. FR-RSA, on the other hand, yielded a strong increase in the RSM correspondence between Model 1 and the target, while the increase was much weaker for Model 2. Please note that a small increase in performance is also expected for the inferior model, since it was designed to exhibit a partial correspondence with ground truth. Overall, this simulation highlights both important aspects of FR-RSA: First, feature reweighting can improve the correspondence between model and target RSMs. Second, FR-RSA is able to recover the ground truth model.

### 3.2. Reweighting units of computational models

#### 3.2.1. Reweighting model units consistently improves correspondence between two RSMs

Our first aim was to evaluate whether FR-RSA reliably increases the correspondence between two RSMs. To this end, we applied FR-RSA to seven different layers of three different DNNs to predict RSMs of behavior, higher visual cortex (HVC) or early visual cortex (EVC), as measured with fMRI in several publicly available datasets (see Datasets and computational models for details). Figure 5 shows the results of comparing classical RSA with FR-RSA across all 168 combinations of analyses. Irrespective of the kind of image set, we found that FR-RSA robustly increased the fit between two RSMs as compared to classical RSA. A chi-squared test revealed that a significantly larger proportion of RSM comparisons showed improvements (144) than comparisons that were worse (24) after reweighting DNN units (*χ*^2^(1, N = 168) = 85.71, *p* < .001). Altogether, in 119 cases (70.83%) the fit between two RSMs was significantly increased when feature reweighting was applied to units of DNN layers. In 8 cases (4.76%) FR-RSA performed significantly worse than classical RSA, while in 41 cases (24.41%) the difference to classical RSA was not significant. Breaking this down into different types of analyses, for the prediction of behavioral RSMs, FR-RSA significantly outperformed classical RSA in 59 cases, with 5 cases that showed significantly worse performance of FR-RSA and 20 cases with a non-significant difference between both methods. Similarly, for the prediction of fMRI RSMs, FR-RSA significantly outperformed classical RSA in 60 cases, performed significantly worse in 3 cases, and showed non-significant differences in 21 cases. Together, these results demonstrate that FR-RSA robustly improves the correspondence between DNNs, brain activity, and behavior.

**Figure 5:**
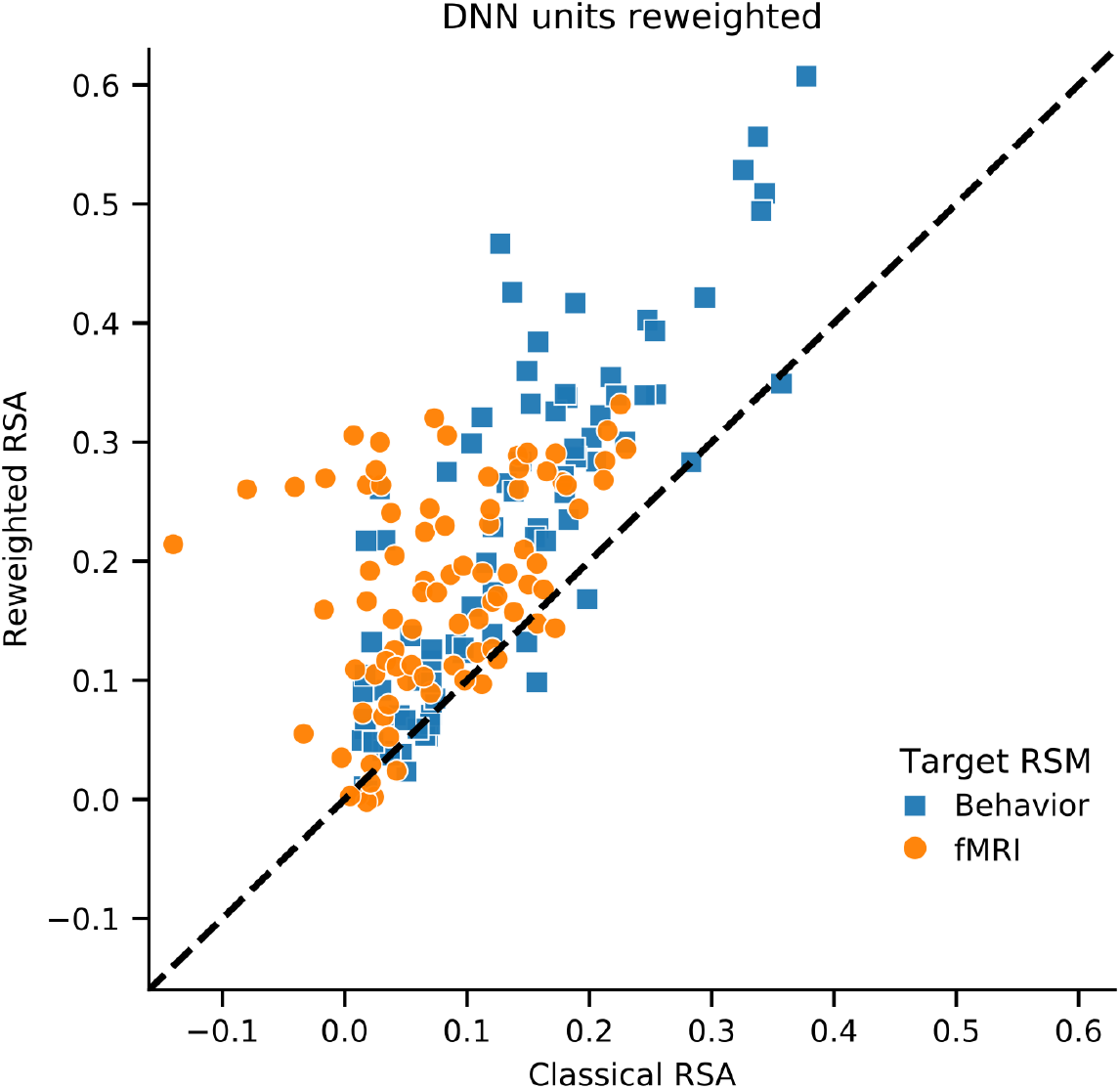
Comparison of classical RSA with feature-reweighted RSA when reweighting DNN units. For most comparisons, feature-reweighted RSA reveals a stronger RSM correspondence than classical RSA (144 stronger, 24 weaker). Of all 168 comparisons, FR-RSA significantly outperformed classical RSA in 119 cases, often leading to strong increases in the fit between two RSMs.

#### 3.2.2. Reweighting model units influences model selection

Having demonstrated the reliable performance of FR-RSA, we next sought to evaluate whether applying FR-RSA also leads to changes in the model selection process: Does the same model produce the best fit to brain or behavioral data, regardless of whether classical RSA or FR-RSA is used, or can FR-RSA lead to qualitative changes in the results, leading to different models that are chosen as optimal (Storrs et al., 2021)? To this end, we assessed the relative predictive performance of three common DNN architectures (AlexNet, VGG-16, ResNet-50) for each of seven layers, when relating their classical and reweighted RDMs to target RSMs derived from either, behavior, HVC, or EVC. These results are shown in Figure 6. In the following we will highlight a subset of all findings.

**Figure 6:**
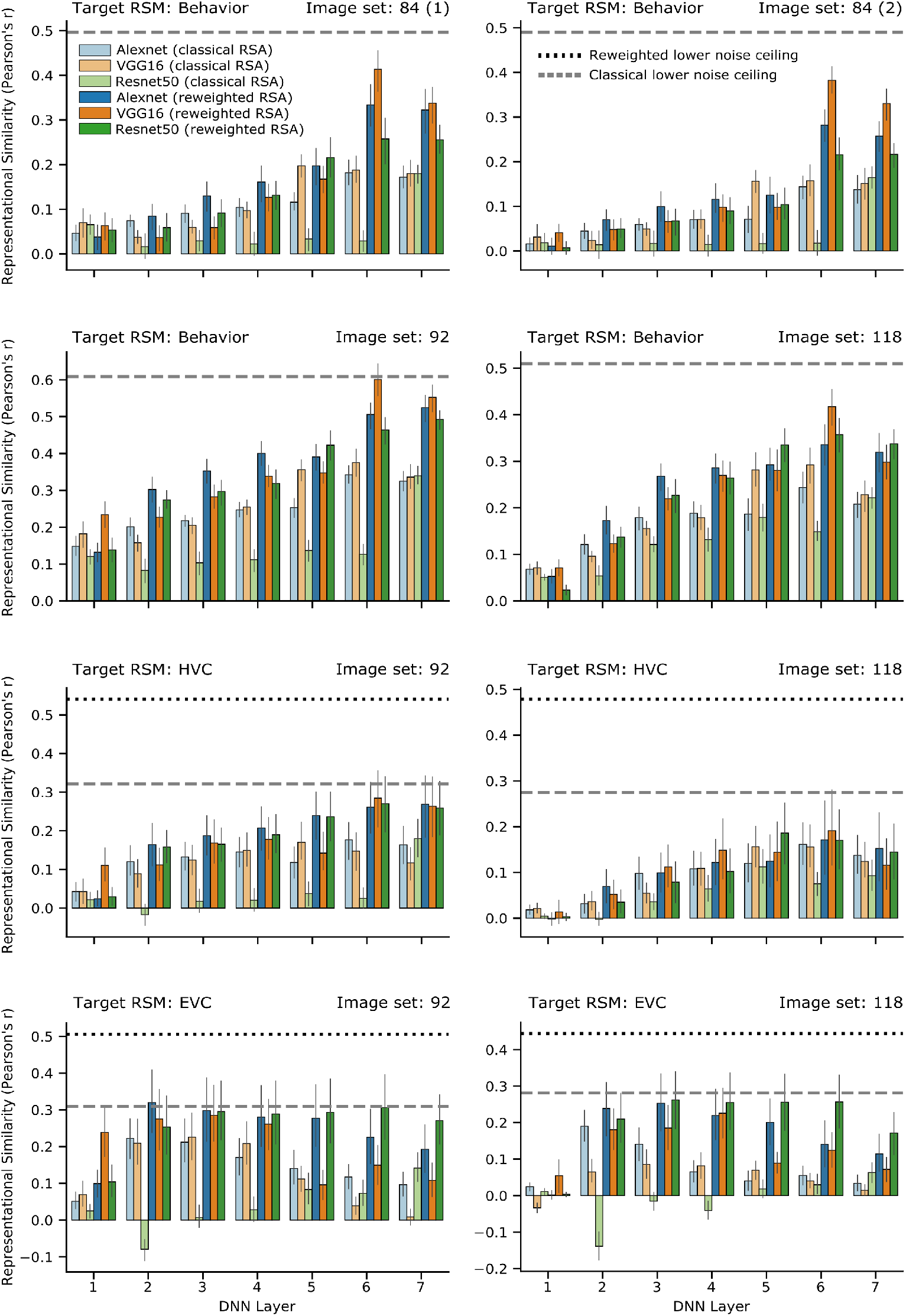
Detailed comparison of classical and feature-reweighted RSA applied to DNN units. Each panel shows the predictive performance of three DNNs (blue: AlexNet, orange: VGG-16, green: ResNet-50) with classical and feature-reweighted RSA (light and dark hues, respectively), for a target RSM of a given image set as indicated above each panel. Target RSMs were derived from either behavior, early (EVC) or higher visual cortex (HVC). In each panel, the dashed gray and the dotted black line indicate the lower classical and lower reweighted noise ceiling, respectively. In many cases, feature reweighting leads to an increased fit between RSMs, as can be seen by the dark bars being generally higher than their light counterparts. In addition, feature reweighting affected model selection, as indicated by the changes in the relative heights of the different model bars for dark and light hues. Note that in cases where behavioral RSMs are the target RSMs, the calculation of reweighted noise ceilings was not possible (see main text). Error bars indicate 95% confidence intervals computed using bootstrapping. Most fits are significantly different from zero.

Let us first focus on the DNN layers’ scores for the image set 118 and EVC target RSM (see Figure 6, bottom right panel). Before reweighting, AlexNet and VGG-16 were correlated significantly with the target RSM across most layers, while ResNet-50 showed fewer significant effects (range of correlations across all layers: AlexNet: 0.024 to 0.191, *M* = 0.079, all layers significant; VGG-16: −0.034 to 0.087, *M* = 0.047, layers 2-6 significant; ResNet-50: −0.14 to 0.063, *M* = −0.011, layers 1, 6, 7 significant; all *p* < 0.035, FDR corrected). AlexNet and VGG-16 also performed significantly better than ResNet-50 for layers 1-6 and 2-5, respectively, while ResNet-50 outperformed AlexNet and VGG-16 only for layer 7 and layers 1 and 7, respectively (all *p* < 0.009, FDR corrected). This picture changed strongly after reweighting DNN units of each layer. ResNet-50’s performance improved strongly (range of correlations: 0.003 to 0.27, *M* = 0.209; layers 2-7 significant; *p* < 0.001, FDR corrected) and no longer showed a significant difference from reweighted AlexNet’s or VGG-16’s performance for layers 1-3 and for layers 1 and 4, respectively. ResNet-50 in fact outperformed AlexNet for layers 4-7 and VGG-16 for layers 2-3 and 5-7, respectively (all *p* < 0.008, FDR corrected). Based on these results, which show that ResNet-50 is the superior model across multiple layers for EVC, it becomes evident that feature-reweighted RSA does, indeed, affect model selection.

The general pattern that FR-RSA selects another model than classical RSA as the best model can be observed when shifting the focus from one specific dataset to all panels in Figure 6. Before feature-reweighting, VGG-16 is very often the best performing model for layer 5 (range of correlations: 0.07 to 0.357; *M* = .19). For 7/8 combinations of image set and target RSM, VGG-16 performed significantly better than both AlexNet and ResNet-50 in layer 5 (all *p* < 0.004, FDR corrected). After feature-reweighting, though, ResNet-50 was the superior model (range of correlations: 0.104 to 0.426; *M* = .263), being significantly better than AlexNet and VGG-16 in 4/8 and 7/8 cases for layer 5 (all *p* < 0.006, FDR corrected). Note that, for the remaining cases, ResNet-50 was never significantly worse than AlexNet or VGG-16. A similar trend can be observed for layer 6 when comparing non-reweighted AlexNet to reweighted VGG-16. These results support the notion that model selection is affected by applying feature-reweighted RSA irrespective of the exact combination of image set and target RSM.

Taken together, there is strong evidence that model selection is influenced by applying feature-reweighted RSA to DNN units. These results highlight the importance of considering alternatives to classical RSA for comparing competing models and suggest the general utility of FR-RSA for adjudicating amongst them.

### 3.3. Voxel-reweighted RSA: Reweighting individual voxels improves prediction of model RSMs, MEG data, and behavior

The second major aim of this study was to explore a novel use case of feature-reweighting: rather than reweighting individual units of a computational model, we tested the degree to which reweighting brain measurements can improve the ability to predict a computational model’s RSM. This approach parallels multivariate decoding, which also reweights individual measurement channels (e.g. voxels) to maximize the fit with a target variable. To this end, we applied FR-RSA to fMRI data from higher and early visual cortex to predict RSMs either from behavior, DNN layers, or MEG time points (MEG-fMRI fusion). The results of all 568 comparisons between classical RSA and voxel-reweighted RSA are shown in Figure 7. FR-RSA applied to voxels overwhelmingly increased the correspondence between various predicting and target RSMs. A chi-squared test revealed that a significantly larger proportion of RSM comparisons showed improvements (370) than comparisons that were worse (198) after reweighting fMRI voxels (χ^2^(1, N = 568) = 52.09, *p* < .001). Overall, in 235 cases (41.37%) the fit between two RSMs increased significantly, in 66 (11.62%) cases FR-RSA performed significantly worse than classical RSA, and in 267 (47.01%) cases the difference to classical RSA was not significant. Again, breaking this down into different types of analyses, for the prediction of behavioral RSMs, FR-RSA significantly outperformed classical RSA in all 4 cases. Similarly, for the prediction of DNN RSMs, FR-RSA significantly outperformed classical RSA in 80 cases, never performed significantly worse, and showed non-significant differences in 4 cases. Finally, when predicting MEG RSMs, FR-RSA significantly outperformed classical RSA in 151 cases, performed significantly worse in 66 cases, and showed non-significant differences in 263 cases. Please note that a large number of the MEG comparisons include MEG samples during which likely no information was present at all. Yet, we chose all time points for a conservative estimate.

**Figure 7:**
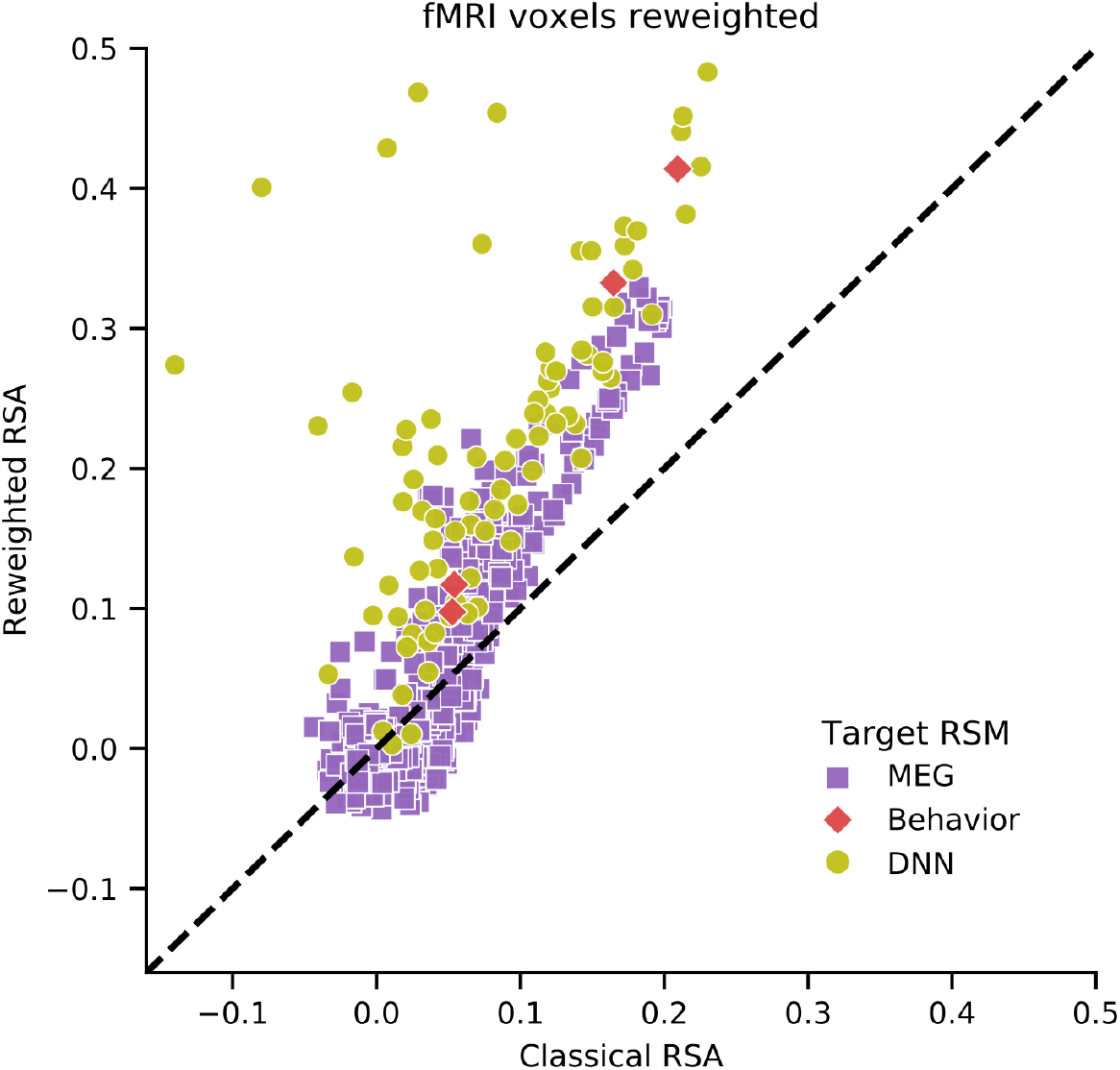
Comparison of classical RSA with feature-reweighted RSA when reweighting fMRI voxels. For most comparisons, feature-reweighted RSA reveals a stronger RSM correspondence than classical RSA when applied to voxels (370 vs. 198). FR-RSA significantly outperformed classical RSA in 235 cases, often leading to very strong increases in the fit between two RSMs.

Focusing on individual results, Figure 8 shows how well reweighted fMRI voxels from two different ROIs explain behavioral RSMs with classical and FR-RSA. While EVC voxels already predicted behavioral RSMs significantly before reweighting (*r* = 0.054 and *r* = 0.053 for image set 92 and 118, respectively), these correlations were increased significantly after reweighting (*r* = 0.117 and *r* = 0.098; *p* < 0.001 FDR corrected for all correlations, *p* < 0.05 uncorrected for the differences). The improvement in RSM correlations was even stronger for HVC: the RSM correspondence before reweighting (r = 0.209 and *r* = 0.165) was again significantly improved after reweighting took place (r = 0.414 and *r* = 0.333; *p* < 0.001 FDR corrected for all correlations, *p* < 0.001 uncorrected for the differences).

**Figure 8:**
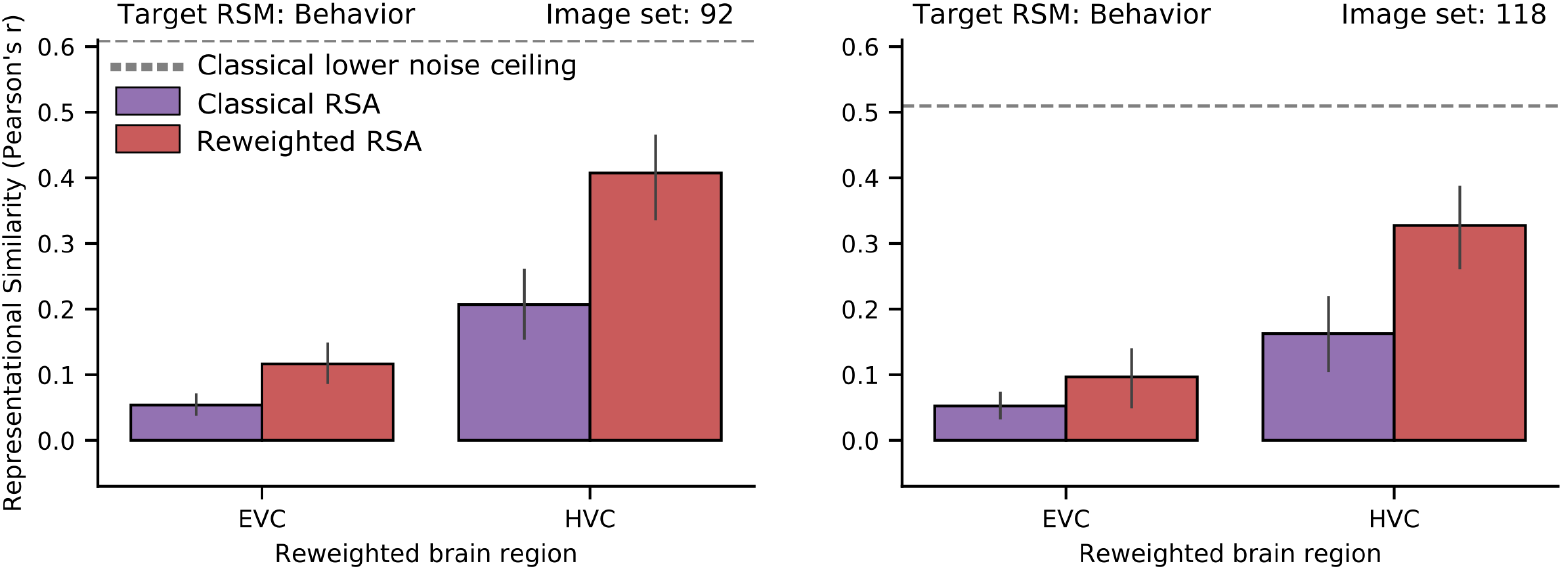
Voxel-reweighted RSA improves the prediction of behavioral similarity from EVC and HVC. Results across both image sets (92, 118) and regions of interest (EVC, HVC) yield consistent increases in representational similarity when applying voxel-reweighted RSA. The dashed gray line indicates the lower classical noise ceiling. The computation of a reweighted noise ceiling was not possible (see main text). Error bars indicate 95% confidence intervals computed using bootstrapping.

Figure 9 shows the relative correspondence of the three DNN architectures for each of seven layers, when relating their RSMs to either classical or reweighted voxel RSMs derived from either HVC or EVC. Overall, reweighting individual voxels led to even stronger improvements in RSM correspondence than reweighting DNN units, at times approaching the reweighted lower noise ceiling. In all four panels in Figure 9, all of the 84 RSM correlations for classical RSA are significantly below the classical lower noise ceiling (all *p* < 0.04, uncorrected). However, for voxel-reweighted RSA, there were seven cases in which a DNN layer’s RSM and brain RSM correlated to an extent that was not significantly different from the *reweighted* lower noise ceiling. These results demonstrate that voxel reweighted RSA can strongly improve the fit between RSMs.

**Figure 9:**
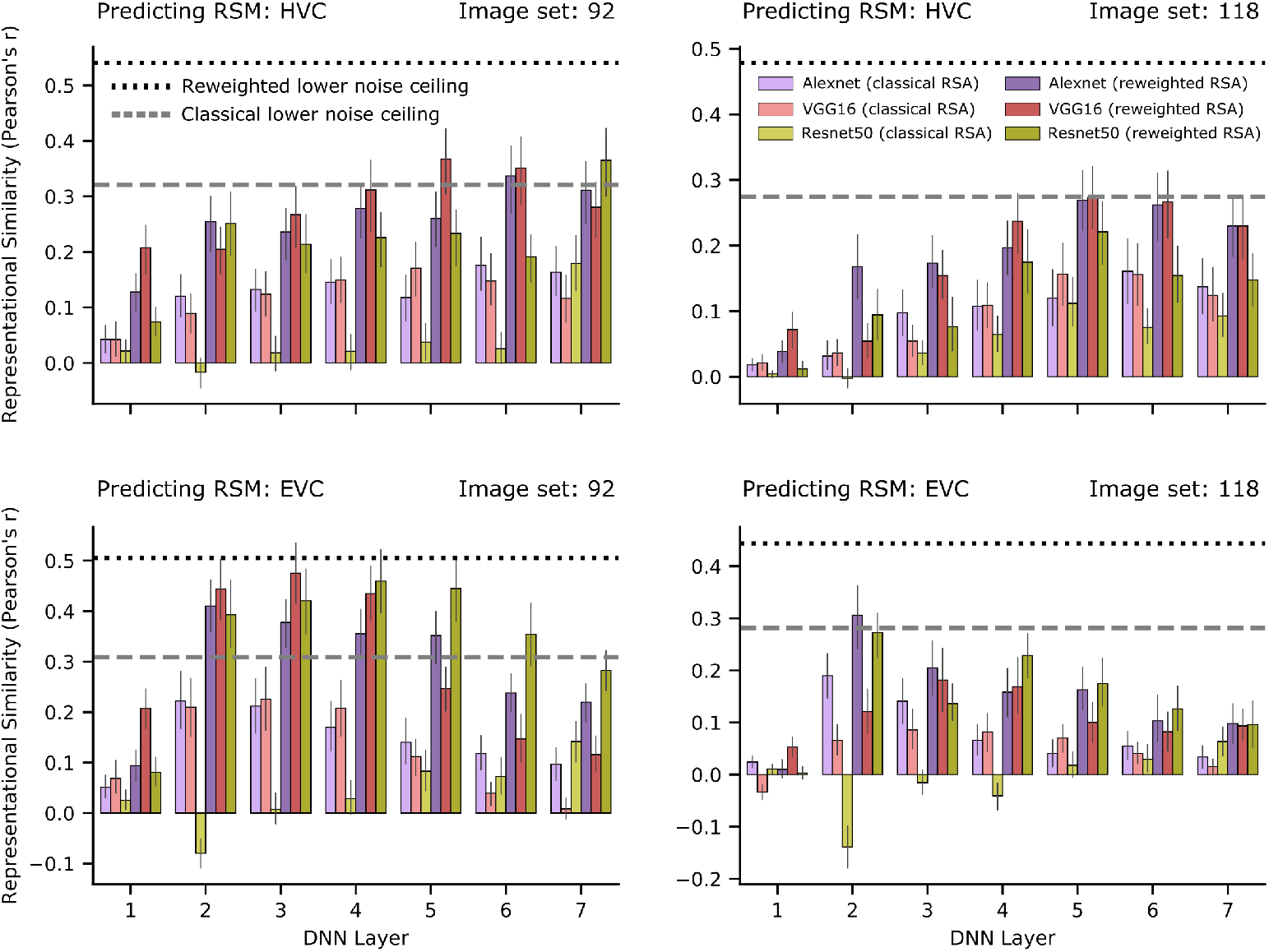
Detailed comparison of classical and voxel-reweighted RSA. Reweighting of individual fMRI voxels leads to strong and consistent increases in the fit between RSMs. Each panel shows how well the RSMs of three DNNs (purple: AlexNet, red: VGG-16, yellow: ResNet-50) can be explained with classical and feature-reweighted RSA (light and dark hues, respectively), when applied to either higher (HVC) or early visual cortex (EVC) RSM of a given image set as indicated above each panel. In each panel, the dashed gray and the dotted black line indicate the lower classical and lower reweighted noise ceiling, respectively. Error bars indicate 95% confidence intervals computed using bootstrapping. Most fits are significantly different from zero.

### 3.4. Reweighting amplifies existing and reveals new peaks when applied to MEG-fMRI fusion

To test whether results generalize beyond the prediction of DNN layer and behavioral RSMs, we conducted MEG-fMRI fusion with classical and feature-reweighted RSA for two datasets for which fMRI and MEG data were available (Figure 10). For each participant separately, we reweighted fMRI voxels at each time point of MEG data, to best predict MEG similarity. For most time points, classical and FR-RSA each yielded a representational similarity significantly larger than zero, as indicated by the purple and red horizontal lines in Figure 10, respectively. Further, for many time points, there were significant differences between classical and FR-RSA (uncorrected), as indicated by the black horizontal line in Figure 10. Overall, FR-RSA revealed peaks which would not have been detected using classical RSA, and it also markedly increased existing peaks.

**Figure 10:**
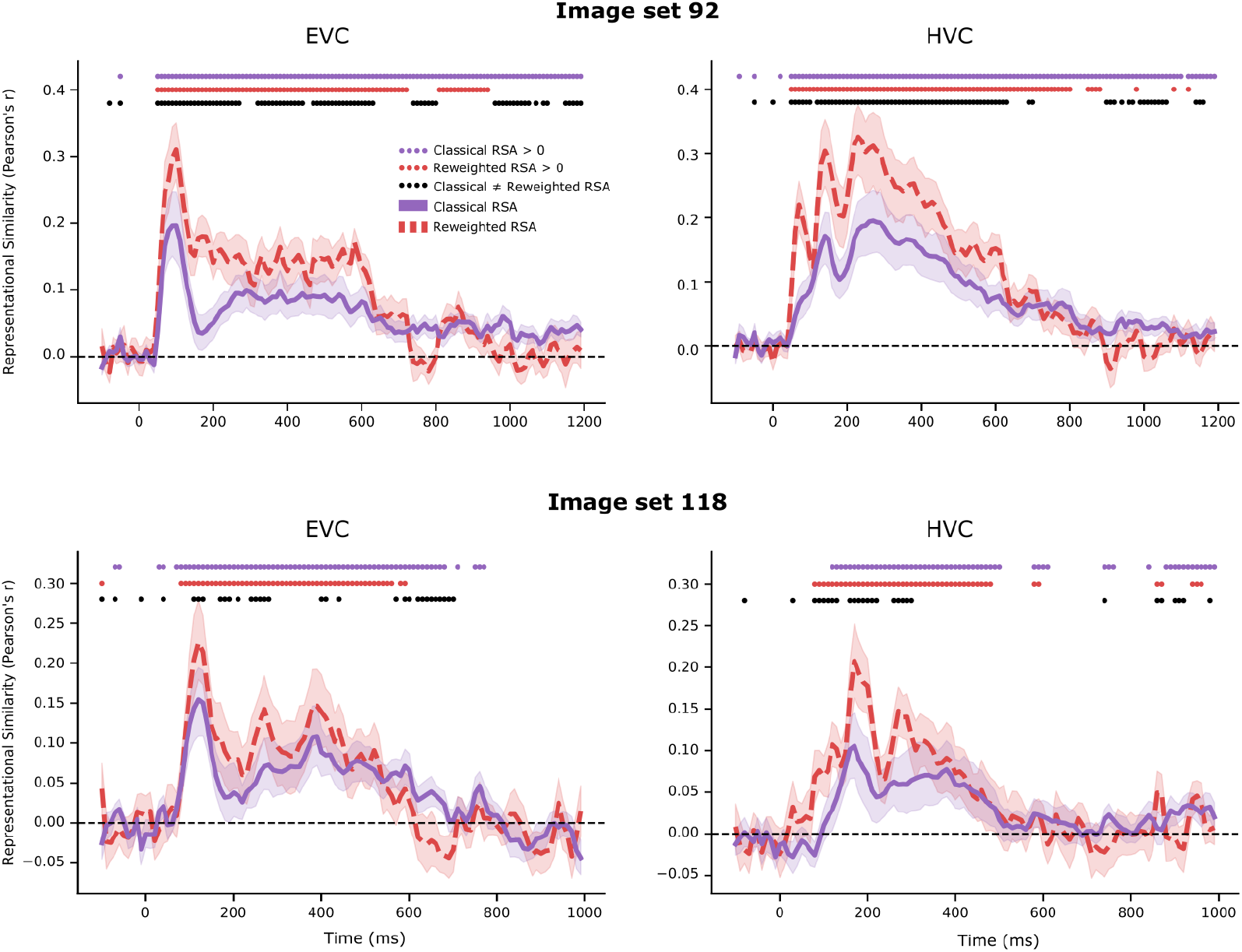
Voxel-reweighted RSA applied to MEG-fMRI fusion. MEG-fMRI fusion can be improved consistently by applying voxel-reweighted RSA, amplifying existing peaks and yielding new ones. Each panel reflects a combination of one image set and brain region, with MEG-fMRI fusion for classical (purple) and feature-reweighed (red) RSA. Purple and red dots indicate time points for which classical and voxel-reweighted RSA yield RSM correlations significantly bigger than zero, respectively. Black dots indicate time points with significant differences between classical and voxel-reweighted RSA (all uncorrected). Shaded areas indicate 95% confidence intervals computed using bootstrapping.

### 3.5. The effect of reweighted noise ceilings on the interpretation of reweighted RSM correspondences

As argued earlier, for feature-reweighted RSA, classical noise ceilings underestimate the best possible performance any model can achieve, invalidating inferences based on classical noise ceilings (see Estimating reweighted noise ceilings). Traditionally, models numerically exceeding the lower noise ceiling are often interpreted as fully explaining the data at hand (Khaligh-Razavi and Kriegeskorte, 2014). To identify whether we find cases where reweighting noise ceilings would change this interpretation of results, we inspected all 736 combinations of analyses, counted the number of cases that numerically exceeded the classical lower noise ceiling, and compared this with the number of cases that numerically exceeded the reweighted lower noise ceiling. Note that of all 736 combinations, only those 168 entered this analysis for which reweighted noise ceilings could be computed. Indeed, a total of 21 reweighting analyses (12.5%) exceeded the classical lower noise ceiling, while no reweighting analysis exceeded the reweighted lower noise ceiling, a significant difference as revealed by a chi-squared test (χ^2^(1, N = 21) = 21, *p* < 0.001). Therefore, following this tradition would lead to an inflated number of results that are reported as fully capturing the data. However, while four analyses even numerically exceeded the classical upper noise ceiling, it is also important to note that none of these results were significant. Thus, while we did not find empirical evidence for results significantly above the classical upper noise ceiling, reweighted noise ceilings are required to prevent biased assessment of a model’s performance.

### 3.6. Does feature reweighting lead to positively biased results?

The analyses presented so far demonstrate that feature-reweighted RSA often increases RSM correspondence and influences model selection. However, it is also necessary to assess whether feature-reweighting leads to positively biased results, meaning above-chance representational similarities despite the absence of any association between representational spaces.

To this end, we carried out a randomization procedure by permuting the condition labels of the target RSM while leaving intact the order of the labels of the predicting RSM. In this case, feature-reweighting should, on average, not lead to an improved RSM correspondence. For a given randomization iteration, we applied the same shuffling to the RSMs of all participants, computed the representational similarity using Pearson’s correlation, and averaged the resulting correlation coefficients, leading to one group mean correlation coefficient for a given permutation. We then repeated this procedure 100 times to generate a distribution of group mean representational similarities for a given combination of predicting and target RSM. This randomization procedure was conducted for four separate such combinations. In all cases, the RSM correspondence yielded a distribution centered around zero (range of means: −0.0003 to 0.0017, see Supplemental Figure S3 for a depiction of empirical null distributions). Together, this showcases that FR-RSA does not lead to positively biased representational similarities.

## 4. Discussion

RSA is widely used to assess the correspondence between brains, behavior, and models and select amongst several candidates the model that best explains a given representational space (Kriegeskorte et al., 2008a; Kriegeskorte and Kievit, 2013). In this work, we evaluated a powerful extension of classical RSA called feature-reweighted RSA (FR-RSA) in which individual features of a predicting RSM are reweighted to maximize the fit with a target RSM. Using fMRI, MEG, and behavioral data from multiple neuroscientific studies as well as several DNNs as computational models, we broadly validated the general applicability of this approach. Further, we present an important novel use case of FR-RSA by applying feature reweighting to brain measurement channels; compared to classical RSA, voxel-reweighted RSA leverages more of the multivariate information content present in human brain (dis-)similarity data, thus nicely complementing existing multivariate decoding techniques. Altogether, we find strong and robust increases in the fit between RSMs. Changes in the model selection process were also often observed when applying feature-reweighting as opposed to classical RSA. Based on these results, we suggest that FR-RSA applied to brain measurement channels could become an important new method to assess the match between representational spaces.

### 4.1. Past developments of reweighted RSA approaches: Similarities and differences

Classical RSA as introduced by Kriegeskorte et al. (2008a) has been studied extensively as a research method. Below, we will briefly outline past developments leading up to our contribution, as well as similarities and differences to our approach. Khaligh-Razavi and Kriegeskorte (2014) were the first to propose reweighting in the form of layer-reweighted RSA, where an entire layer of a computational model (in this case a DNN) receives a single weight to predict a target RDM. Peterson et al. (2016) were the first to use feature-reweighted RSA (FR-RSA), by reweighting individual DNN units of a fully-connected layer using ridge regression to predict behavioral RSMs. Jozwik et al. (2017) applied a similar approach to predict human object similarity judgments from entire feature maps of convolutional DNN layers or individual units of fully-connected layers. Finally, Storrs et al. (2021) recently proposed a two-stage RSA approach applied to RDMs of human inferior temporal cortex, first reweighting principal components of DNN layers and then combining individual layers together with another reweighting step. When comparing multiple trained DNNs with each other, regarding how well they predict inferior temporal cortex activity after reweighting, they found that performance differences between DNNs were strongly diminished.

While these studies each contributed important novel information and already highlighted the potential value of FR-RSA, our study (1) broadly and systematically validates FR-RSA across numerous behavioral and neuroimaging datasets, thus confirming the value of FR-RSA beyond individual previous studies, (2) provides a new use case of FR-RSA by applying reweighting to individual voxels, offering a powerful new method for assessing the fit of brain data with models and behavior, and (3) introduces feature-reweighted noise ceilings, providing a more suitable approach for evaluating the upper limit of the predictive performance of any model given the available data.

While all previous FR-RSA approaches have in common the reweighting of individual features, there are also important differences. Several previous approaches (Jozwik et al., 2016, 2017; Storrs et al., 2021) have limited themselves to non-negative weights given that true dissimilarities can only be positive, while our proposed FR-RSA approach avoids this constraint. As a consequence, our approach not only stretches or squeezes individual features, but can also invert their values. If a researcher assumes that each feature of the model in question should be regarded as a distinct property (like color or orientation) that corresponds to a separate similarity matrix, weights should be non-negative to avoid illegal feature-specific (dis-)similarities. If, though, one regards features not as distinct properties, but merely as features, we believe that, given the potential information contained in individual units, it makes sense to allow weights to take on any value to possibly counteract contributions of otherwise overrepresented features. However, given the previous use of the non-negativity constraint, we repeated a subset of analyses with such a constraint and found results to be similar (see Supplemental Figure S2). Further, our approach additionally uses an L2 penalty similar to Peterson et al. (2016) and Jozwik et al. (2017), but in contrast to Jozwik et al. (2016) and Storrs et al. (2021). This choice of regularization, however, is reasonable given the expected collinearity of features. While Storrs et al. (2021) countered multicollinearity with principal components regression, our choice of ridge regression provides smoother shrinkage of regression parameters and may lead to slightly improved prediction (Hastie et al., 2009). Finally, during cross-validation, Peterson et al. (2016) left out individual object pair similarities, while we and others (e.g. Jozwik et al., 2017) left out entire objects, thus avoiding potential leakage effects given that object pair similarities are not all independent. While Storrs et al. (2021) cross-validated across both participants and stimuli and used bootstrapping for estimating statistical significance, our approach of cross-validating across stimuli alone leaves the option to carry out reweighting at the participant level, thus allowing classical statistical analyses for inferring that the effect found in each participant is present in the population. Beyond being computationally more efficient, FR-RSA at the participant level offers an approach with well-known statistical properties for the generalization to the population , which may be more challenging for double cross-validation that mixes the sources of variance for objects and participants.

### 4.2. A novel approach for feature reweighting

Further, different from all previous developments, we present a novel approach for feature reweighting, by applying it to brain activity patterns: voxel-reweighted RSA. This application was motivated by classical multivariate decoding. In multivariate decoding, individual voxels receive their own weights which reflect their importance in optimally conducting a linear read-out of a binary (e.g. stimulus category) or a continuous target variable (e.g. stimulus size). However, in the context of RSA, where all voxels receive equal weights, this approach to our knowledge has not been applied previously. Thus, relative to multivariate decoding, classical RSA may underestimate the linear information content of multivariate measurements. By reweighting individual voxels to optimally predict (dis-)similarity, feature-reweighted RSA can leverage more of the rich multivariate information content of the data.

### 4.3. Reweighted noise ceilings for reweighted RSA approaches

The fit between a model and data is limited by the quality of the model and the noise in the data. Noise ceilings provide an estimate of the best performance any model can achieve and thus allow us to tell how far off a given model is, given the noise in the data. However, when reweighting individual features, classical noise ceilings underestimate this upper performance threshold, since they themselves do not take reweighting into account. This can lead to a situation in which reweighted RSMs may exceed the classical lower noise ceiling but not the reweighted lower noise ceiling, thus leading to falsely interpreting the reweighted RSM to explain the data at hand when only considering classical noise ceilings, a result we confirmed empirically. In the context of feature reweighting, we therefore suggest calculating *reweighted* noise ceilings, which again provide a sensible performance corridor for reweighted RSMs. Note that previous adaptations regarding the calculation of noise ceilings address different problems, such as how to calculate noise ceilings for a model that received weights when fitted to a single group-average target RDM (or RSM) (Storrs et al., 2020).

### 4.4. Use cases for feature-reweighted RSA

While FR-RSA generally yielded strong improvements of the correspondence between computational models and brain data and also affected which model was selected among competing models, it may be argued that, while feature reweighting is computationally feasible, it should not be applied to model RSMs in general (see Storrs et al., 2021, for a previous discussion of this topic). According to this line of reasoning, the representational similarity between a model and a given dataset already provides a good estimate of the explanatory power of this model, and reweighting the model’s features would be akin to testing the performance of a different model. To illustrate this line of reasoning, assume for a moment that a model RSM is built not from computational models but originates from an experimental design with several factors. For example, in an experiment, participants may have been presented with images of faces with different degrees of 3D rotation, which can be quantified by the three parameters pitch, roll, and yaw. Each of the three orientation directions would thus constitute a feature. A model RSM in this experiment could simply quantify the similarity of face orientation between faces integrated across all features. When a researcher is interested primarily in the fit of such a static model, we would argue that reweighting of individual model features should not be applied since it would change what hypothesis is tested. However, if each model feature is treated as a separate variable of interest, for which the contribution to a target RSM is unknown, then reweighting can improve the fit, and indeed, this approach is already commonly used in practice when conducting RSA in a multiple regression framework (e.g. Jozwik et al., 2016; Groen et al., 2018; Hebart and Baker, 2018).

Likewise, for computational models, when the model is treated as a good approximation of all relevant aspects of a brain or behavioral dataset, we would argue that reweighting should not be carried out. For example, a learned DNN model can, among others, be characterized as a product of its architecture (e.g. number of layers and units per layer, transition functions, etc.), its learning objective (e.g. object classification), and the stimuli and object classes that had been used during training (Kietzmann et al., 2019; Richards et al., 2019). When testing the degree to which all of these aspects are already representative of brain and behavioral datasets, applying reweighting may distort this assessment. However, it is well known that commonly used datasets for training DNNs do not reflect the categories most relevant to humans (Hebart et al., 2019; Mehrer et al., 2021), and that the learning objective of ventral visual cortex is known to go beyond simple object classification (Kravitz et al., 2013). Thus, successful feature reweighting promises to yield a better match beyond the images a DNN had been trained on and beyond its limited training objective, possibly better reflecting the explanatory power of a DNN architecture trained on object images. Likewise, in principle, the reweighting can even be reversed in the DNN weights between layers, yielding a better match to the target RSM without affecting model performance. More generally, when interested merely in the information contained in a given computational model, we would argue that FR-RSA can be applied more liberally. Thus, whether FR-RSA should be applied to features of a computational model depends entirely on what aspects of the computational model are supposed to be fixed and what aspects are allowed to vary. Crucially, researchers should be explicit about this choice in their studies to avoid confusion and draw valid conclusions.

When it comes to reweighting of measurement channels (e.g. voxel-reweighted RSA), the applicability of this approach again depends on the aims of the researcher and their assumption about the nature of the representations studied. When interested in testing the existence of representational similarity alone (i.e. “Does the model show *any* fit to activity patterns in brain region X?”), which is a very common goal for RSA, we argue that reweighting of measurement channels (e.g. voxel-reweighted RSA) can be carried out more generally. Drawing the parallel to multivariate decoding, voxel-reweighted RSA would allow weighting individual voxels in a way that reflects a plausible lower bound of the potential representations that can be read-out by downstream regions (Kriegeskorte and Bandettini, 2007). Whether these representations are indeed used by other brain regions or behavior remains an empirical question for both multivariate decoding and RSA (Williams et al., 2007; Ritchie et al., 2019). Given its increased statistical power, applying FR-RSA to measurement channels promises to advance our understanding of representational content in a way similar to how multivariate decoding has leveraged information contained in measured brain activity patterns.

However, when using RSA for carrying out model comparisons (i.e. “Which model best explains activity patterns in brain region X?”), there are certain restrictions to the use of voxel reweighting. Assume for the moment that we are dealing with the ventral visual cortex as a region of interest and that this region includes face-selective clusters (e.g. fusiform face area, FFA). Ventral visual cortex is known to represent objects in a distributed fashion, while FFA responds more uniformly to images of faces. When comparing a simpler model that tests for face selectivity alone against a more complex model testing for object selectivity including faces, the simpler model may win over the more complex one simply because feature reweighting may focus on face selective voxels for the simpler model, which may be easier to fit than the more complex model that is based on representations with more distributed voxel activity patterns. Thus, for model comparisons, voxel-reweighted RSA would not be testing the degree to which an entire region is well-suited for characterizing a model but may focus on selective parts of these regions, which may even be different for each model. In such a case, finding a subspace of the predicting RSM that best fits the target RSM does not imply that this subspace is (or should be) reflective of the entire target. This may, of course, be a desirable side effect of FR-RSA, and, indeed, the feature weights could be inspected to test the degree to which this is the case. However, if one would like to treat a region as carrying a more-or-less homogeneously and widely-distributed representation, then voxel-reweighted RSA may complicate model comparisons.

Based on these considerations, if a researcher has the theoretical possibility to reweight either, model units or fMRI voxels, we argue that, for model comparison, it is the model units that ought to be reweighted. If model comparison is not of interest and the primary aim is assessing whether a model and brain RSM show any correspondence, we argue that one may default to voxel reweighting.

### 4.5. Possible extensions of feature-reweighted RSA

There are several ways to refine feature-reweighting in the context of RSA. First, other penalization regimes could be applied. Instead of using an L2-penalty, one could either use an L1-penalty or a combination of L1- and L2-penalization (or pruning, see Tarigopula et al., 2021). We opted for the L2-penalization since we did not want to select a subset of features (as any penalization regime utilizing the L1-norm would do) and since an L1-penalization would incur a greater computational load. Second, one could not only penalize the predictors’ variances but also their covariance so that all features that exhibit a high covariance with other features are penalized. This approach might, however, strongly increase the computational load of the fitting procedure. A third possible extension of feature-reweighting would be to fit weights bidirectionally. That way, both RSMs would receive weights to optimally predict the other RSM, possibly using a latent vector approach (e.g. canonical correlation analysis). Fourth, the fitting procedure could be repeated so that the residuals of the first fitting procedure are predicted by a linear weighted combination of some other predicting RSM. Finally, a feature-reweighting that automatically selects the best reweighting options from those just mentioned could be combined with reweighting entire layers of a DNN (i.e. two-stage RSA, Storrs et al., 2021). The bottleneck for implementing such a procedure will be computational limits with regards to CPU and RAM resources and the complexity of cross-validation schemes for identifying hyperparameters and splitting data in independent folds.

### 4.6. Considerations when using FR-RSA

In addition to broadly validating feature-reweighting and exploring a novel use case of it, we also provide an implementation of FR-RSA in Python (https://github.com/ViCCo-Group/frrsa). In the following, we would like to provide important considerations when using FR-RSA and mention possible drawbacks.

The first aspect to consider is that FR-RSA utilizes cross-validation to prevent overfitting and nested cross-validation to identify the optimal regularization parameter for ridge regression. Both outer and inner crossvalidation require data to be split into independent training and test sets. Please note that the cross-validation was performed across images and not runs as is common in multivariate decoding. For the outer cross-validation, by default, FR-RSA uses 5-fold cross-validation, repeated ten times with different random splits. On a subset of the analyzed data (i.e. for 36 different combinations of predicting RSM, target RSM, and image set), we assessed different fold sizes of the outer cross-validation post-hoc and found results to be largely unaffected (see Supplemental Figure S4). We opted for 5-fold cross-validation because it provides enough data for stable estimation of the statistical models in the training set, while at the same time not consuming too many computational resources for actually fitting the models. This crossvalidation was repeated ten times to make sure that many different object pairs will at some point be part of training or test folds. For the inner crossvalidation, 5-fold cross-validation is used and repeated five times with different random splits. We tested different numbers of repetition post-hoc and found that results were largely unaffected by how the inner cross-validation was set up (see Supplemental Figure S5). We opted to repeat the inner cross-validation five times as a good balance between how well the best hyperparameter for a given outer cross-validation is estimated (more repetitions should lead to a better estimation) and computational load. Note that increasing either the fold size or the number of repetitions can have noticeable effects on the computation resources needed to run the algorithm.

Further, a question researchers who want to deploy FR-RSA might have is what is the minimum number of conditions FR-RSA requires to be used successfully and how FR-RSA performance scales with this number. On a subset of the analyzed data (i.e. for 120 different combinations of predicting RSM, target RSM, and image set), we repeatedly subsampled from all available conditions (in our case images) and assessed how FR-RSA performed in comparison to classical RSA. We found that, on average, FR-RSA almost always performed better than classical RSA, with the performance of FR-RSA increasing with the number of drawn images (see Supplemental Figure S6). The results indicate that FR-RSA can be used successfully with a comparably small number of conditions but benefits from more conditions.

A drawback of FR-RSA, in comparison to classical RSA, is the higher computational load, specifically for models with a large number of features, such as early layers of a DNN. For many different computational problem sizes without non-negativity constraint on the *β* weights, we measured how much time and RAM were needed to solve the problem when the *β* weights were allowed to take on any value (see Supplemental Figures S7 and S8). Note, however, that computational resources are much higher when imposing a non-negativity constraint and that the resources needed might also depend on the hardware of the machine in question, the operating system that machine uses, and other software-specific factors.

### 4.7. Conclusion

Representational Similarity Analysis (RSA) has emerged as a popular tool for relating representational spaces of the brain, computational models, and behavior to each other (Kriegeskorte et al., 2008a). As such, it can reveal which model best captures how the brain represents relations between stimuli. Feature-reweighted RSA, the approach we investigated here, not only consistently increases the fit between RSMs, but also affects which models are best at reproducing a given brain’s representational geometry. Further, when applied not to model units but to brain measurement channels, voxel-reweighted RSA more fully leverages the information content present in representational spaces of the brain and thus nicely complements classical multivariate decoding. Overall, FR-RSA is well suited to become a general-purpose method for measuring the information content shared between representations in computational models, brain, and behavior, and may improve our ability as scientists to adjudicate between competing models.

## Supporting information

complete supplemental material

## Acknowledgments

The authors would like to thank Katherine R. Storrs, Tim C. Kietzmann and Nikolaus Kriegeskorte for helpful discussions, Jacob L. S. Bellmund, Felix Deilmann, Magdalena Gippert and Joshua Grant for useful comments on an earlier version of the manuscript, and Hannes Hannsen for improvement of the code base.

## References

Bankson, B., Hebart, M., Groen, I., Baker, C., 2018. The temporal evolution of conceptual object representations revealed through models of behavior, semantics and deep neural networks. NeuroImage 178, 172–182. URL: https://linkinghub.elsevier.com/retrieve/pii/S1053811918304440, doi:10.1016/j.neuroimage.2018.05.037.

Benjamini, Y., Hochberg, Y., 1995. Controlling the false discovery rate: A practical and powerful approach to multiple testing. Journal of the Royal Statistical Society: Series B (Methodological) 57, 289–300. URL: https://rss.onlinelibrary.wiley.com/doi/abs/10.1111/j.2517-6161.1995.tb02031.x.

Bobadilla-Suarez, S., Ahlheim, C., Mehrotra, A., Panos, A., Love, B.C., 2020. Measures of neural similarity. Computational Brain & Behavior 3, 369–383. URL: http://link.springer.com/10.1007/s42113-019-00068-5, doi:10.1007/s42113-019-00068-5.

Charest, I., Kriegeskorte, N., Kay, K.N., 2018. GLMdenoise improves multivariate pattern analysis of fMRI data. NeuroImage 183, 606–616. URL: https://www.sciencedirect.com/science/article/pii/S1053811918307626, doi:10.1016/j.neuroimage.2018.08.064.

Cichy, R.M., Khosla, A., Pantazis, D., Torralba, A., Oliva, A., 2016. Comparison of deep neural networks to spatio-temporal cortical dynamics of human visual object recognition reveals hierarchical correspondence. Scientific Reports 6, 27755. URL: http://www.nature.com/articles/srep27755, doi:10.1038/srep27755.

Cichy, R.M., Kriegeskorte, N., Jozwik, K.M., van den Bosch, J.J., Charest, I., 2019. The spatiotemporal neural dynamics underlying perceived similarity for real-world objects. NeuroImage 194, 12–24. URL: https://www.sciencedirect.com/science/article/pii/S1053811919302083, doi:10.1016/j.neuroimage.2019.03.031.

Cichy, R.M., Pantazis, D., Oliva, A., 2014. Resolving human object recognition in space and time. Nature Neuroscience 17, 455–462. URL: http://www.nature.com/articles/nn.3635, doi:10.1038/nn.3635.

Diedrichsen, J., Kriegeskorte, N., 2017. Representational models: A common framework for understanding encoding, pattern-component, and representational-similarity analysis. PLOS Computational Biology 13, e1005508. doi:10.1371/journal.pcbi.1005508.

Diedrichsen, J., Yokoi, A., Arbuckle, S.A., 2018. Pattern component modeling: A flexible approach for understanding the representational structure of brain activity patterns. NeuroImage 180, 119–133. URL: https://www.sciencedirect.com/science/article/pii/S1053811917306985, doi:10.1016/j.neuroimage.2017.08.051.

Glasser, M.F., Coalson, T.S., Robinson, E.C., Hacker, C.D., Harwell, J., Yacoub, E., Ugurbil, K., Andersson, J., Beckmann, C.F., Jenkinson, M., Smith, S.M., Van Essen, D.C., 2016. A multi-modal parcellation of human cerebral cortex. Nature 536, 171–178. URL: http://www.nature.com/articles/nature18933, doi:10.1038/nature18933.

Groen, I.I.A., Greene, M.R., Baldassano, C., Fei-Fei, L., Beck, D.M., Baker, C.I., 2018. Distinct contributions of functional and deep neural network features to representational similarity of scenes in human brain and behavior. eLife, e32962 doi:10.7554/eLife.32962.

Hastie, T., Tibshirani, R., Friedman, J.H., 2009. The elements of statistical learning: Data mining, inference, and prediction. 2 ed., Springer.

Haxby, J.V., Connolly, A.C., Guntupalli, J.S., 2014. Decoding neural representational spaces using multivariate pattern analysis. Annual Review of Neuroscience 37, 435–456. URL: http://www.annualreviews.org/doi/10.1146/annurev-neuro-062012-170325, doi:10.1146/annurev-neuro-062012-170325.

Haynes, J.D., Rees, G., 2006. Decoding mental states from brain activity in humans. Nature Reviews Neuroscience 7, 523–534. URL: http://www.nature.com/articles/nrn1931, doi:10.1038/nrn1931.

He, K., Zhang, X., Ren, S., Sun, J., 2016. Deep residual learning for image recognition, in: 2016 IEEE Conference on Computer Vision and Pattern Recognition (CVPR), pp. 770–778. URL: http://ieeexplore.ieee.org/document/7780459/, doi:10.1109/CVPR.2016.90.

Hebart, M.N., Baker, C.I., 2018. Deconstructing multivariate decoding for the study of brain function. NeuroImage 180, 4–18. URL: https://www.sciencedirect.com/science/article/pii/S1053811917306523, doi:10.1016/j.neuroimage.2017.08.005.

Hebart, M.N., Dickter, A.H., Kidder, A., Kwok, W.Y., Corriveau, A., Van Wicklin, C., Baker, C.I., 2019. THINGS: A database of 1,854 object concepts and more than 26,000 naturalistic object images. PLOS ONE 14, e0223792. doi:10.1371/journal.pone.0223792.

Hoerl, A.E., Kennard, R.W., 1970. Ridge regression: Biased estimation for nonorthogonal problems. Technometrics 12, 55–67. URL: https://www.tandfonline.com/doi/abs/10.1080/00401706.1970.10488634, doi:10.1080/00401706.1970.10488634.

Hout, M.C., Goldinger, S.D., Ferguson, R.W., 2013. The versatility of SpAM: A fast, efficient, spatial method of data collection for multidimensional scaling. Journal of Experimental Psychology: General 142, 256–281. URL: https://psycnet.apa.org/record/2012-17536-001, doi:10.1037/a0028860.

Jozwik, K.M., Kriegeskorte, N., Mur, M., 2016. Visual features as stepping stones toward semantics: Explaining object similarity in it and perception with non-negative least squares. Neuropsychologia 83, 201–226. URL: http://www.sciencedirect.com/science/article/pii/S0028393215301998, doi:10.1016/j.neuropsychologia.2015.10.023.

Jozwik, K.M., Kriegeskorte, N., Storrs, K.R., Mur, M., 2017. Deep convolutional neural networks outperform feature-based but not categorical models in explaining object similarity judgments. Frontiers in Psychology 8, 1726. URL: https://www.frontiersin.org/article/10.3389/fpsyg.2017.01726, doi:10.3389/fpsyg.2017.01726.

Khaligh-Razavi, S.M., Kriegeskorte, N., 2014. Deep supervised, but not unsupervised, models may explain it cortical representation. PLoS Computational Biology 10, e1003915. doi:10.1371/journal.pcbi.1003915.

Kietzmann, T.C., McClure, P., Kriegeskorte, N., 2019. Deep neural networks in computational neuroscience. Oxford Research Encyclopedia of Neuroscience URL: https://oxfordre.com/neuroscience/view/10.1093/acrefore/9780190264086.001.0001/acrefore-9780190264086-e-46, doi:10.1093/acrefore/9780190264086.013.46.

Kravitz, D.J., Saleem, K.S., Baker, C.I., Ungerleider, L.G., Mishkin, M., 2013. The ventral visual pathway: an expanded neural framework for the processing of object quality. Trends in Cognitive Sciences 17, 26–49. URL: https://www.sciencedirect.com/science/article/pii/S1364661312002471, doi:10.1016/j.tics.2012.10.011.

Kriegeskorte, N., Bandettini, P., 2007. Analyzing for information, not activation, to exploit high-resolution fMRI. NeuroImage 38, 649–662. URL: https://www.sciencedirect.com/science/article/pii/S1053811907001188, doi:10.1016/j.neuroimage.2007.02.022.

Kriegeskorte, N., Kievit, R.A., 2013. Representational geometry: Integrating cognition, computation, and the brain. Trends in Cognitive Sciences 17. URL: https://www.cell.com/trends/cognitive-sciences/fulltext/S1364-6613(13)00127-7, doi:10.1016/j.tics.2013.06.007.

Kriegeskorte, N., Mur, M., 2012. Inverse MDS: Inferring dissimilarity structure from multiple item arrangements. Frontiers in Psychology 3, 245. URL: https://www.frontiersin.org/article/10.3389/fpsyg.2012.00245, doi:10.3389/fpsyg.2012.00245.

Kriegeskorte, N., Mur, M., Bandettini, P., 2008a. Representational similarity analysis - connecting the branches of systems neuroscience. Frontiers in Systems Neuroscience 2, 4. URL: https://www.frontiersin.org/article/10.3389/neuro.06.004.2008, doi:10.3389/neuro.06.004.2008.

Kriegeskorte, N., Mur, M., Ruff, D.A., Kiani, R., Bodurka, J., Esteky, H., Tanaka, K., Bandettini, P.A., 2008b. Matching categorical object representations in inferior temporal cortex of man and monkey. Neuron 60, 1126–1141. doi:10.1016/j.neuron.2008.10.043.

Krizhevsky, A., Sutskever, I., Hinton, G.E., 2012. ImageNet classification with deep convolutional neural networks, in: Advances in neural information processing systems, p. 1097–1105.

Mehrer, J., Spoerer, C.J., Jones, E.C., Kriegeskorte, N., Kietzmann, T.C., 2021. An ecologically motivated image dataset for deep learning yields better models of human vision. Proceedings of the National Academy of Sciences 118. URL: https://www.pnas.org/content/118/8/e2011417118, doi:10.1073/pnas.2011417118.

Mur, M., Meys, M., Bodurka, J., Goebel, R., Bandettini, P., Kriegeskorte, N., 2013. Human object-similarity judgments reflect and transcend the primate-IT object representation. Frontiers in Psychology 4, 128. URL: https://www.frontiersin.org/article/10.3389/fpsyg.2013.00128, doi:10.3389/fpsyg.2013.00128.

Nili, H., Wingfield, C., Walther, A., Su, L., Marslen-Wilson, W., Kriegeskorte, N., 2014. A toolbox for representational similarity analysis. PLoS Computational Biology 10, 1–11. doi:10.1371/journal.pcbi.1003553.

Peterson, J.C., Abbott, J.T., Griffiths, T.L., 2016. Adapting deep network features to capture psychological representations. arXiv URL: http://arxiv.org/abs/1608.02164.

Ramírez, F.M., Revsine, C., Merriam, E.P., 2020. What do across-subject analyses really tell us about neural coding? Neuropsychologia 143, 107489. URL: https://www.sciencedirect.com/science/article/pii/S0028393220301603, doi:10.1016/j.neuropsychologia.2020.107489.

Richards, B.A., Lillicrap, T.P., Beaudoin, P., Bengio, Y., Bogacz, R., Christensen, A., Clopath, C., Costa, R.P., de Berker, A., Ganguli, S., Gillon, C.J., Hafner, D., Kepecs, A., Kriegeskorte, N., Latham, P., Lindsay, G.W., Miller, K.D., Naud, R., Pack, C.C., Poirazi, P., Roelfsema, P., Sacramento, J., Saxe, A., Scellier, B., Schapiro, A.C., Senn, W., Wayne, G., Yamins, D., Zenke, F., Zylberberg, J., Therien, D., Kording, K.P., 2019. A deep learning framework for neuroscience. Nature Neuroscience 22, 1761–1770. URL: https://doi.org/10.1038/s41593-019-0520-2, doi:10.1038/s41593-019-0520-2.

Ritchie, J.B., Kaplan, D.M., Klein, C., 2019. Decoding the brain: Neural representation and the limits of multivariate pattern analysis in cognitive neuroscience. The British Journal for the Philosophy of Science 70, 581–607. doi:10.1093/bjps/axx023.

Rokem, A., Kay, K., 2020. Fractional ridge regression: a fast, interpretable reparameterization of ridge regression. GigaScience 9. doi:10.1093/gigascience/giaa133.

Russakovsky, O., Deng, J., Su, H., Krause, J., Satheesh, S., Ma, S., Huang, Z., Karpathy, A., Khosla, A., Bernstein, M., Berg, A.C., Fei-Fei, L., 2015. ImageNet large scale visual recognition challenge. International Journal of Computer Vision 115, 211–252. URL: https://doi.org/10.1007/s11263-015-0816-y, doi:10.1007/s11263-015-0816-y.

Simonyan, K., Zisserman, A., 2015. Very deep convolutional networks for large-scale image recognition. arXiv URL: https://arxiv.org/abs/1409.1556, arXiv:1409.1556.

Storrs, K.R., Khaligh-Razavi, S.M., Kriegeskorte, N., 2020. Noise ceiling on the crossvalidated performance of reweighted models of representational dissimilarity: Addendum to khaligh-razavi kriegeskorte (2014). bioRxiv URL: https://www.biorxiv.org/content/10.1101/2020.03.23.003046v1, doi:10.1101/2020.03.23.003046.

Storrs, K.R., Kietzmann, T.C., Walther, A., Mehrer, J., Kriegeskorte, N., 2021. Diverse deep neural networks all predict human inferior temporal cortex well, after training and fitting. Journal of Cognitive Neuroscience 33, 2044–2064. doi:10.1162/jocn\_a\_01755.

Tarigopula, H.P., Fairhall, S.L., Hasson, U., 2021. Improved prediction of behavioral and neural similarity spaces using pruned DNNs. bioRxiv URL: https://www.biorxiv.org/content/10.1101/2021.07.08.451521v1, doi:10.1101/2021.07.08.451521.

Vedaldi, A., Lenc, K., 2015. MatConvNet: Convolutional neural networks for matlab, in: Proceedings of the 23rd ACM International Conference on Multimedia, Association for Computing Machinery, New York, NY, USA. p. 689–692. URL: https://doi.org/10.1145/2733373.2807412, doi:10.1145/2733373.2807412.

Walther, A., Nili, H., Ejaz, N., Alink, A., Kriegeskorte, N., Diedrichsen, J., 2016. Reliability of dissimilarity measures for multi-voxel pattern analysis. NeuroImage 137, 188–200. URL: https://www.sciencedirect.com/science/article/pii/S1053811915011258, doi:https://doi.org/10.1016/j.neuroimage.2015.12.012.

Williams, M.A., Dang, S., Kanwisher, N.G., 2007. Only some spatial patterns of fmri response are read out in task performance. Nature Neuroscience 10, 685–686. URL: https://www.nature.com/articles/nn1900, doi:10.1038/nn1900.

