## Supplementary material for "Feature-reweighted representational similarity analysis: A method for improving the fit between computational models, brains, and behavior": complete supplemental material

Kaniuth, P. & Hebart M. N.

**S1. Effect of z-scoring object-patterns.**

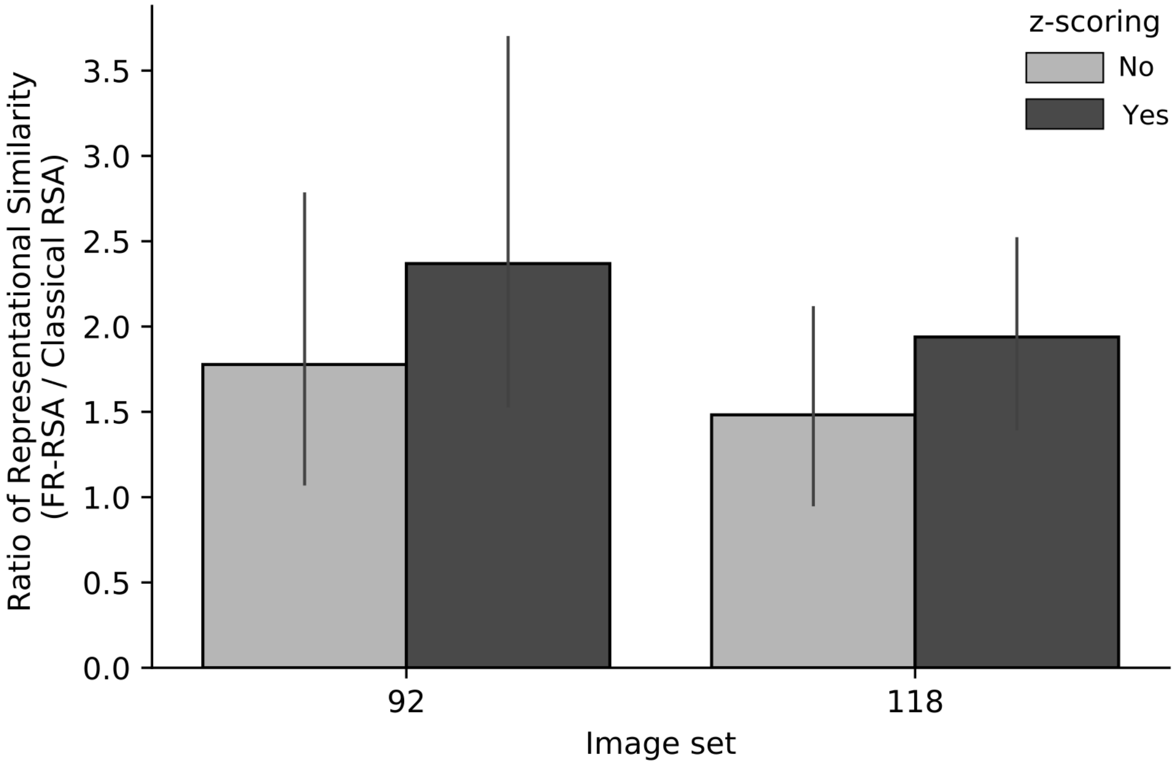


**Supplemental Figure S1.** For a subset of the analyzed data, we post-hoc assessed how z-scoring of the conditions' activity patterns changed the effect of feature-reweighting and found results to be slightly better when z-scoring was applied.

### S2. Effect of non-negativity constraint.

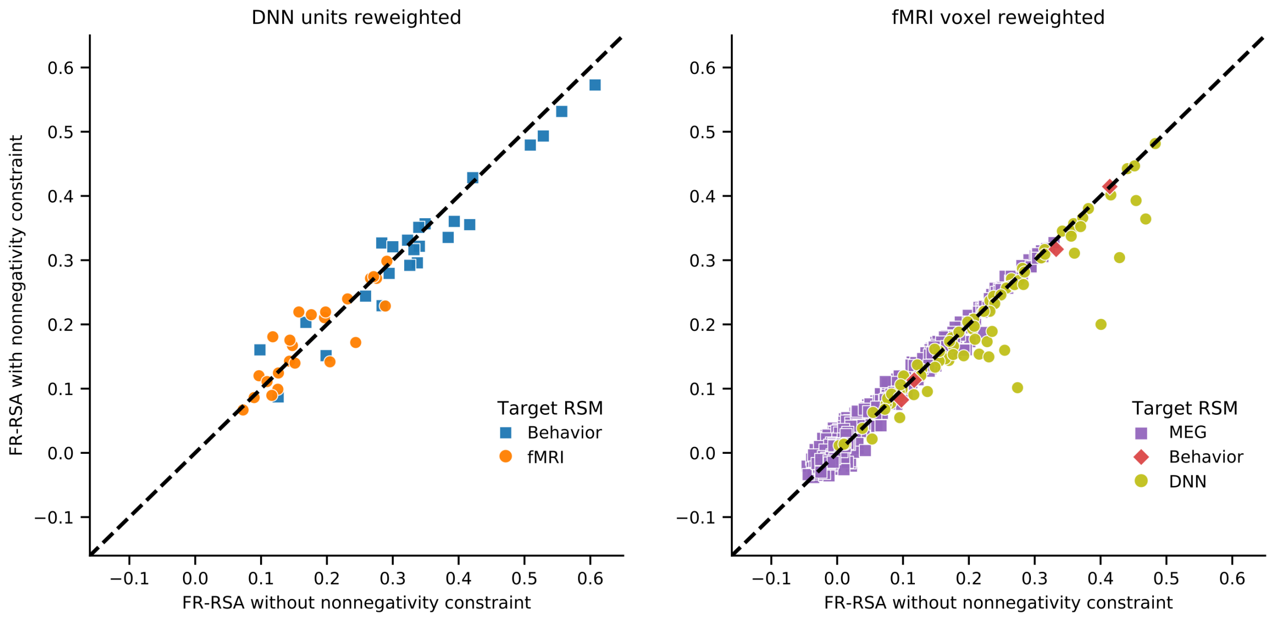


**Supplemental Figure S2.** We post-hoc assessed how imposing a non-negativity constraint on the β weights affects the performance of FR-RSA, for a subset of analyses involving reweighting of DNN units (left) and for all analyses involving reweighting of fMRI voxels (right). Overall, the results were very similar.

### S3. Does FR-RSA lead to positively-biased results?

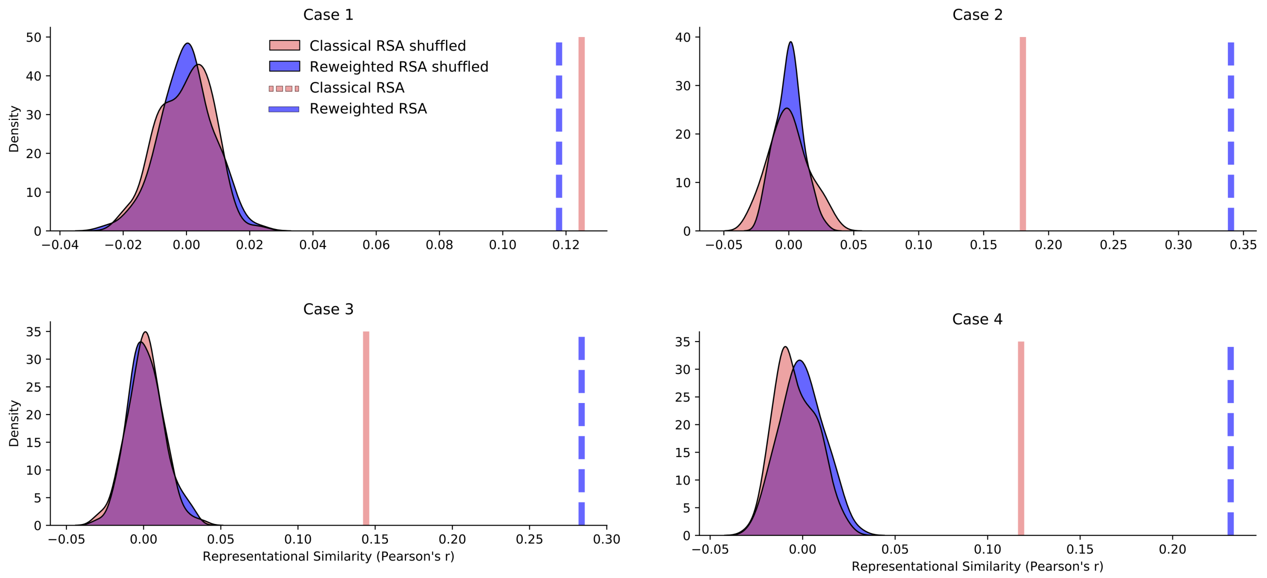


**Supplemental Figure S3.** Feature-reweighting does not produce positively-biased results when applied to a target RSM with shuffled labels. For four test cases (Case 1: image set 118, predicting RSM VGG16's layer 7, target RSM HVC; Case 2: image set 84 (1), predicting RSM VGG16's layer 7, target RSM behavior; Case 3: image set 84 (2), predicting RSM Alexnet's layer 6, target RSM behavior; Case 4: image set 92, predicting RSM Alexnet's layer 6, target RSM EVC) we applied classical and feature-reweighted RSA when randomly shuffling the condition labels of the target RSM, 100 times each. Each panel shows the distribution (using kernel density estimation) of the individual shuffling runs' group-mean scores for classical and feature-reweighted RSA (red and blue, respectively) as well as the group-mean score for classical RSA (red) and feature-reweighted RSA (blue) when applied to the same target RSM with intact labels. In all of the four test cases, for all shuffling runs, the RSM correspondence that resulted from reweighting a target RSM with shuffled labels did not differ significantly from zero and was significantly different from the RSM correspondence that resulted from applying feature-reweighting without shuffling the target RSM's condition labels.

**S4. Effects of different fold sizes of the outer cross-validation**

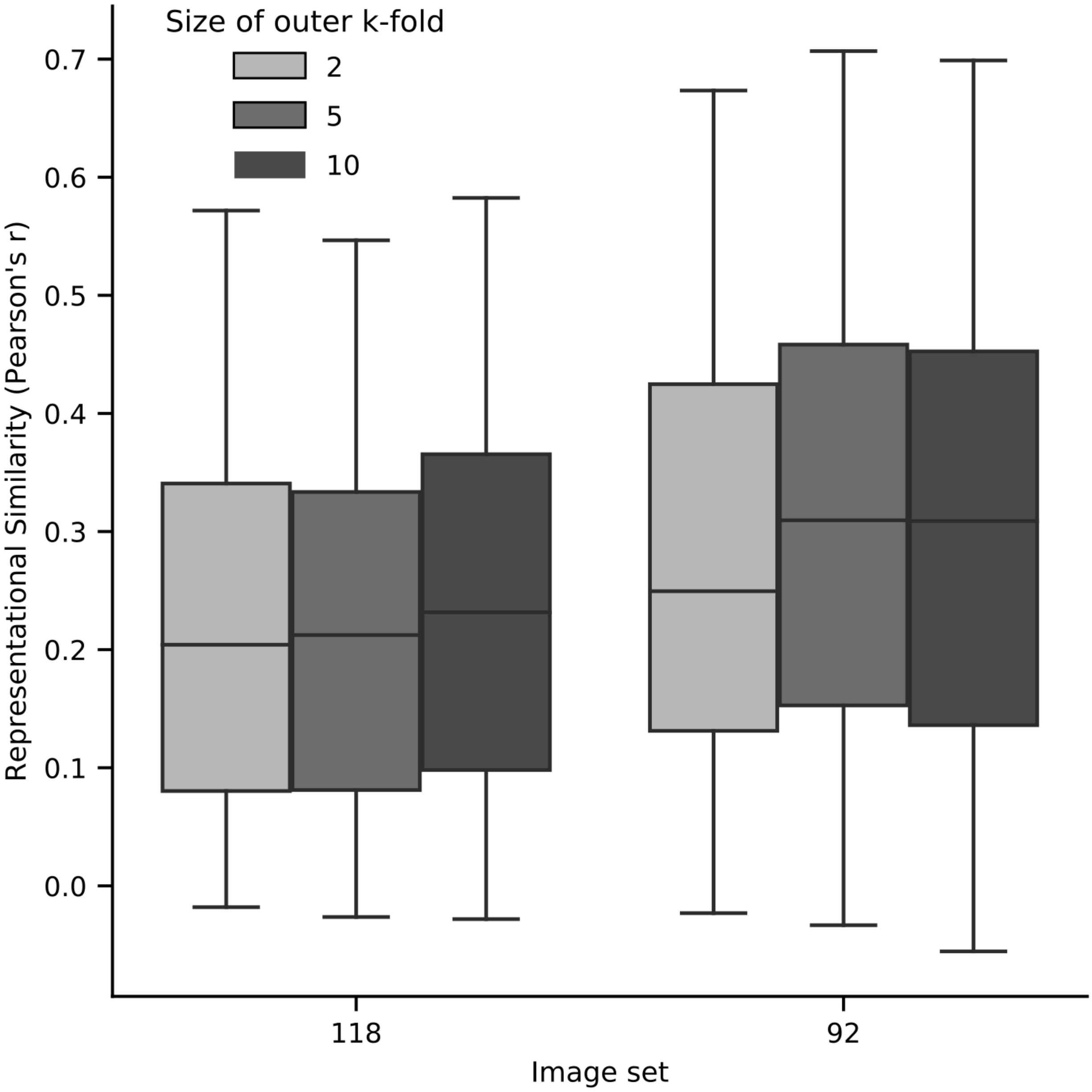


**Supplemental Figure S4.** For a subset of the analyzed data, we systematically varied the size of the outer k-fold in post-hoc analyses. Results were largely invariant across different fold sizes. For computational reasons, we opted for a 5-fold cross-validation, which was repeated ten times.

**S5. Effects of different number of repetitions of the inner cross-validation.**

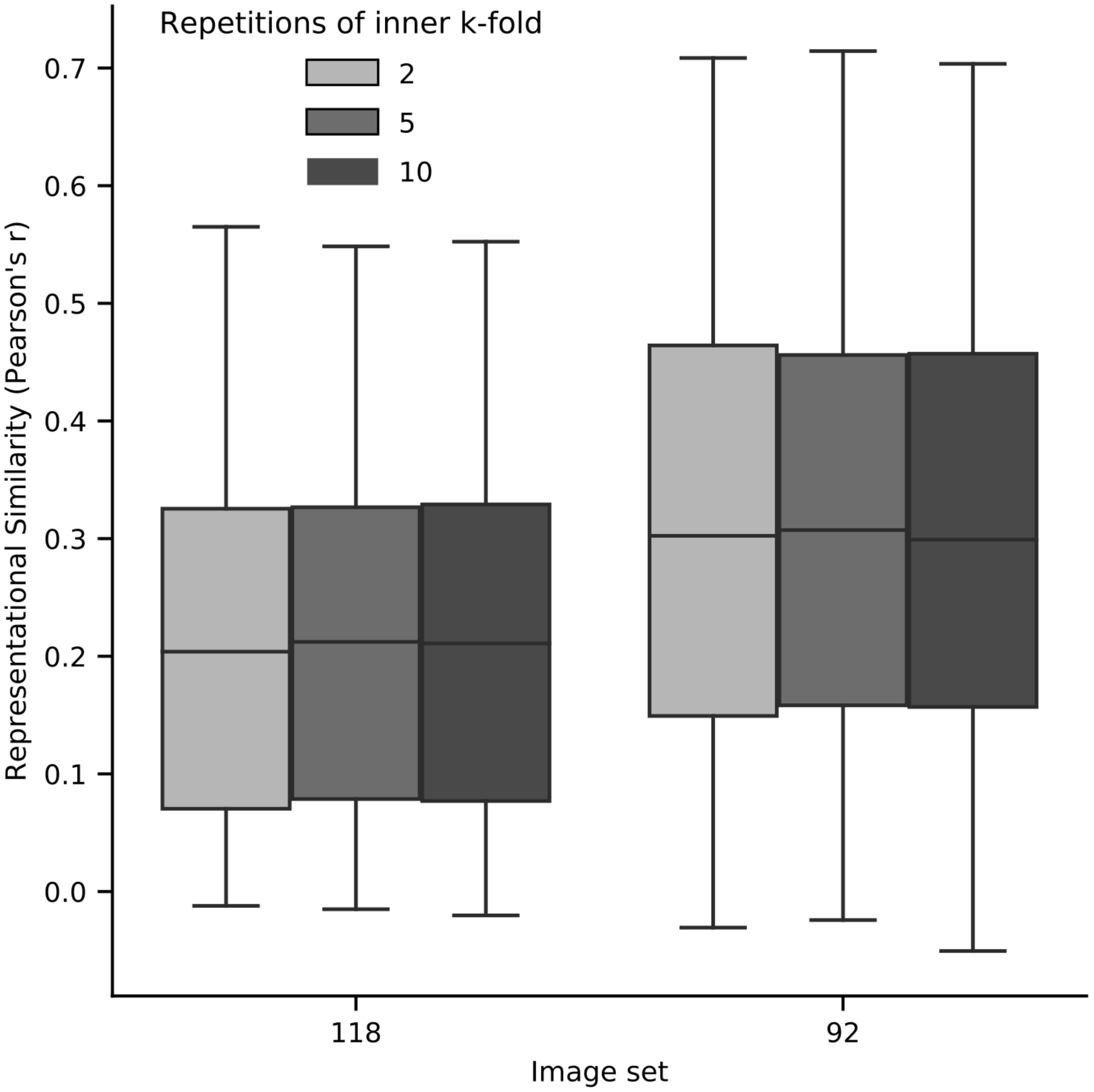


**Supplemental Figure S5.** For a subset of the analyzed data, we systematically varied how often the inner 5-fold cross-validation was repeated. Results were largely invariant across different repetition times. For computational reasons, we opted for repeating the inner cross-validation five times.

### S6. Performance of FR-RSA for repeatedly sampling subsets of conditions.

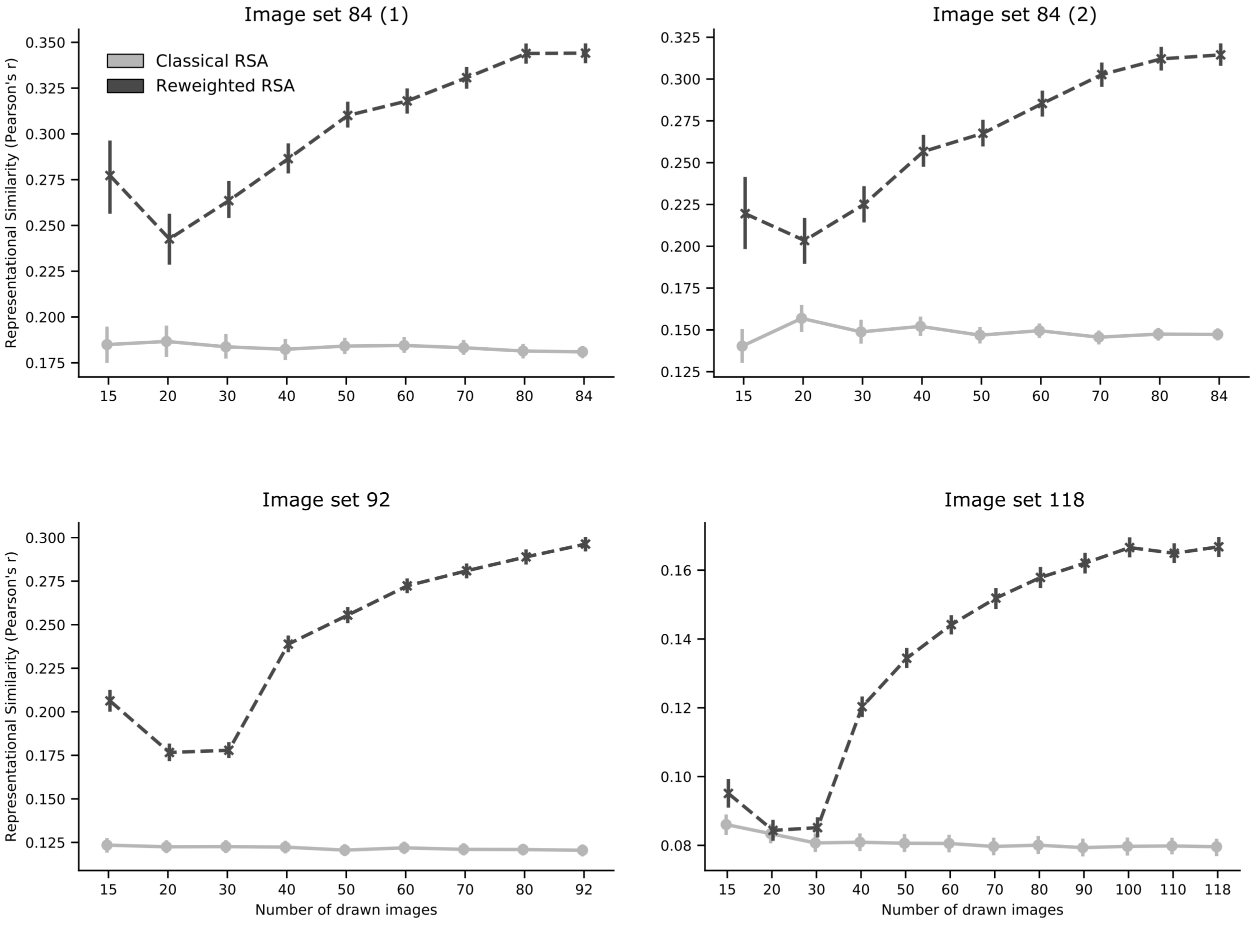


**Supplemental Figure S6.** Comparison of FR-RSA with classical RSA across different numbers of drawn conditions (in our case images) for a range of datasets and models. Number of conditions were subsampled from the original data. Each subsample size was drawn 100 times. Note that in each panel the scores of different combinations of predicting and target RSMs were averaged for a given number of drawn images. Error bars reflect 95% confidence intervals across subsamples computed using bootstrapping.

### S7. Time usage of different computational problems

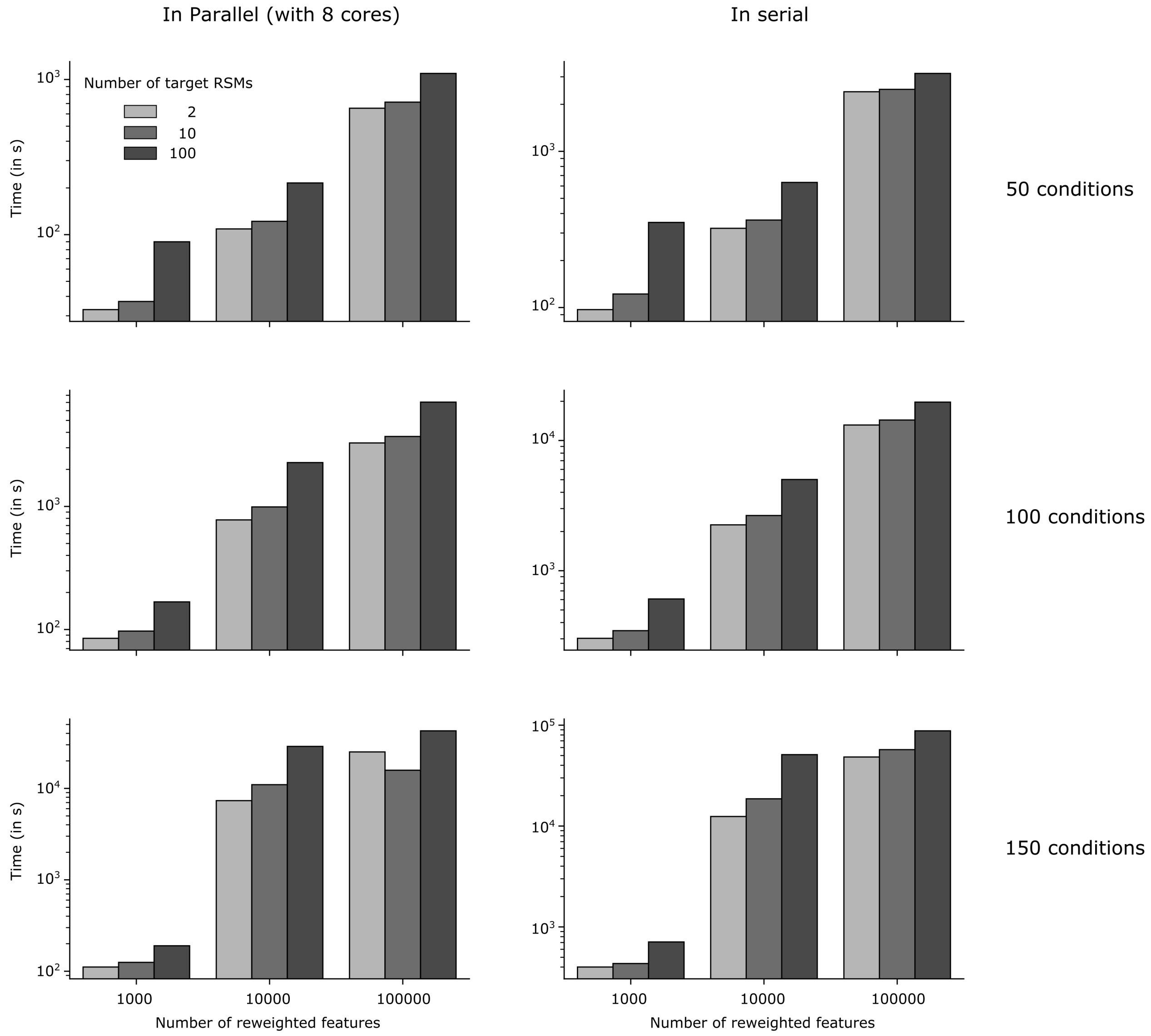


**Supplemental Figure S7.** For different computational problems, we measured how much time was needed to solve the problem. For an outer 5-fold which was repeated ten times, we measured the time in seconds needed (y-axis in log scale with base 10) as a function of the number of reweighted features (x-axis) for different numbers of conditions (rows) and whether the outer cross-validation was explicitly parallelized or not (columns). All analyses were conducted without a no non-negativity constraint on the β weights. Note that the scale of the y-axis differs for each panel.

### S8. RAM usage of different computational problems

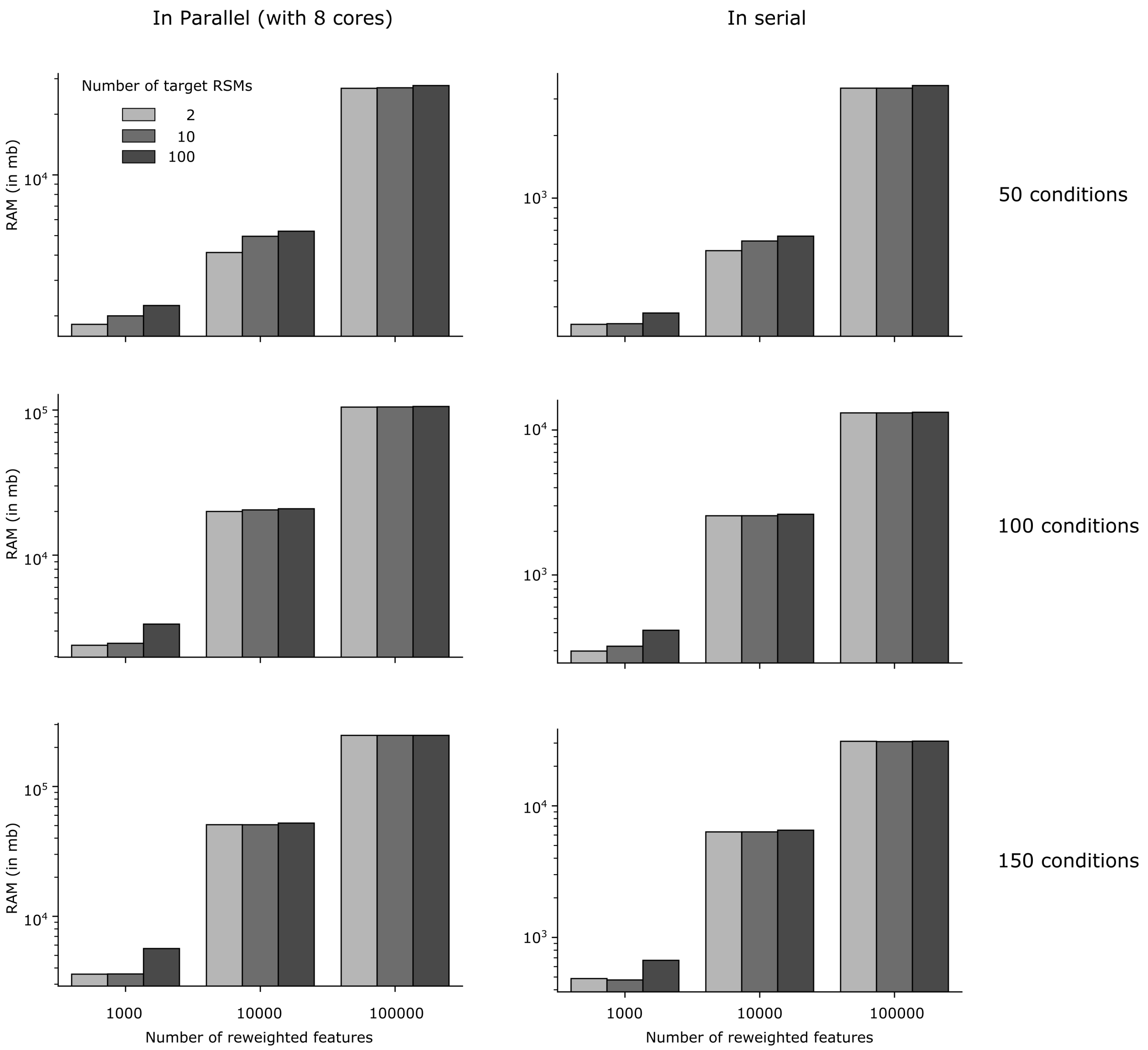


**Supplemental Figure S8.** For different computational problems, we measured how much RAM was needed to solve the problem. For an outer 5-fold which was repeated ten times, we measured the RAM usage in megabyte needed (y-axis in log scale with base 10) as a function of the number of reweighted features (x-axis) for different numbers of conditions (rows) and whether the outer cross-validation was explicitly parallelized or not (columns). All analyses were conducted without a no non-negativity constraint on the β weights. Note that the scale of the y-axis differs for each panel.
